# Histone marks enable formation of immiscible phase-separated chromatin compartments

**DOI:** 10.1101/2023.12.02.569699

**Authors:** Wenjing Sun, Jiacheng Wang, Wenqian Liu, Baixue Tang, Hongyuan Qu, Jianwei Wang, Haitao Li, Wei Xie, Pilong Li

**Affiliations:** State Key Laboratory of Membrane Biology, Beijing Frontier Research Center for Biological Structure, School of Life Sciences, Tsinghua University; Tsinghua-Peking Center for Life Sciences, Beijing 100084, China; Center for Stem Cell Biology and Regenerative Medicine, MOE Key Laboratory of Bioinformatics, School of Life Sciences, Tsinghua University; Tsinghua-Peking Center for Life Sciences, Beijing 100084, China; MOE Key Laboratory of Protein Sciences, Beijing Frontier Research Center for Biological Structure, Department of Basic Medical Sciences, School of Medicine, Tsinghua University, Beijing 100084, China; School of Pharmaceutical Sciences, Tsinghua University, Beijing 100084, China

## Abstract

Eukaryotic chromatin is organized into compartments for gene expression regulation, but the underlying mechanisms remain unclear. Here, we demonstrate that multivalent H3K27me3 and its reader, CBX7-PRC1 complex, regulate facultative heterochromatin via a phase separation mechanism, similar to constitutive heterochromatin^1^. Facultative and constitutive heterochromatin represent distinct, coexisting condensates in nuclei. *In vitro,* H3K27me3- and H3K9me3-marked nucleosomal arrays and their reader complexes can phase separate into immiscible condensates that are analogous to the relationship between facultative and constitutive heterochromatin *in vivo*. Moreover, overexpression of CBX7-PRC1 causes aberrant chromatin compartmentalization *in vivo* as demonstrated by H3K9me3 CUT&Tag and up-regulation of genes related to cancer such as acute myeloblastic leukemia (AML). CBX7 inhibitor effectively inhibits cancer cell proliferation possibly through phase separation-mediated compartment reorganization. Our data demonstrated how the specificity of compartmentalization is achieved based on the formation of immiscible phase-separated condensates, and offer novel epigenetic mechanistic insights into tumor development.

---

Chromatin compartmentalization plays an important role in genome stability and transcriptional regulation^2,3^, in which proteins with related functions accumulate, have been described as biomolecular condensates formed via liquid-liquid phase separation (LLPS). Distinct compartments are enriched with specific histone post-translational modifications (PTMs), which are recognized by their cognate “readers”, often through multivalent interactions^4,5^. Facultative heterochromatic regions are enriched with H3K27me3. H3K27me3 can be recognized by its reader, Polycomb Repressive Complex 1 (PRC1), for chromatin compaction and gene expression suppression^6^. This letter tackles how the specificity of phase-separated compartmentalization is achieved, a frequently asked question in the field of biomolecular condensate. We reveal that chromatin compartmentalization by histone mark-mediated formation of immiscible phase-separated condensates is a critical force in genome organization. Essentially the high degree of cooperativity associated with switch-like phase separation enables a clear distinction between “seemingly promiscuous” biochemical systems, i.e. systems with subtle differences in microscopic interactions. To this end, we focused on the facultative heterochromatin mark, H3K27me3, together with PRC1. The canonical PRC1, also called CBX-PRC1, is composed of four core subunits, RING1a/b, PCGFx (BMI1, MEL18, etc.), PHC1/2/3, and CBX2/4/6/7/8 (ref. ^7^). To determine which CBX protein(s) primarily targets H3K27me3 in NIH-3T3 cells, we examined the colocalization of H3K27me3 with CBXs using immunofluorescence staining. Interestingly, CBX7 shows the highest degree of colocalization with H3K27me3 (Fig. 1a,b), followed by CBX8, CBX6, CBX4, and CBX2 (Fig. 1c,e and Extended Data Fig. 1a). These results suggest that CBX7/8/6 preferentially recognize H3K27me3 in fibroblast. Other subunits of PRC1, such as RING1B and BMI1, also highly colocalize with H3K27me3 (Fig. 1c-e). These results indicate that CBX7-PRC1 is a representative H3K27me3 reader in the facultative heterochromatin compartment in NIH-3T3 cells.

**Fig. 1:**
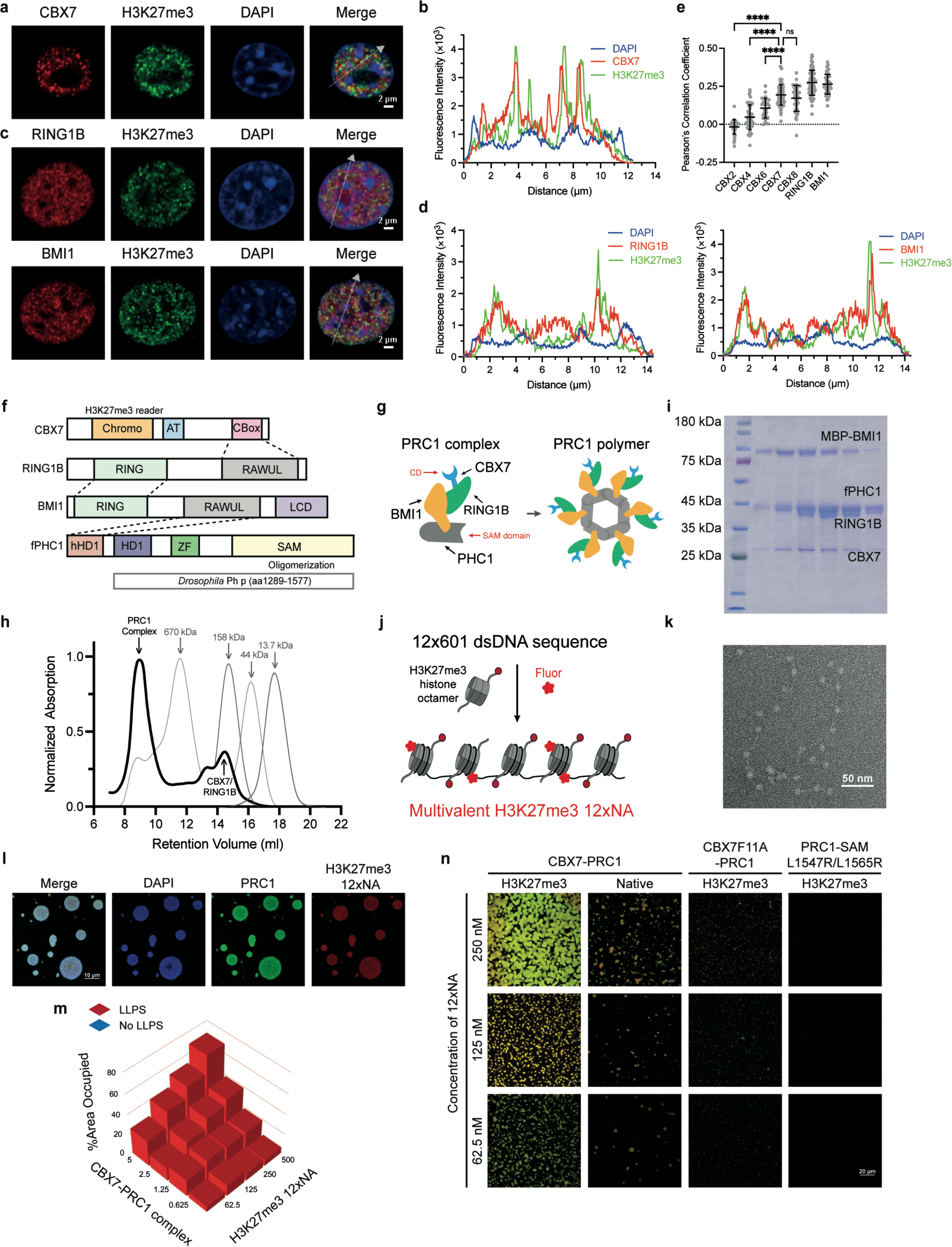
CBX7 and other canonical PRC1 subunits colocalize with H3K27me3 *in vivo* and *in vitro*. **a**, Immunofluorescence images of NIH-3T3 cells show that CBX7 (red) colocalizes with H3K27me3 (green) in facultative heterochromatin. DNA was counterstained with DAPI (blue). **b**, Line scans of the merged image of the cell in (**a**). The fluorescence intensity of CBX7, H3K27me3, and DNA (DAPI) was scanned along the white arrow. **c**, Immunofluorescence images of NIH-3T3 cells show that RING1B (red) (top row), and BMI1 (red) (bottom row), colocalize with H3K27me3 (green) in facultative heterochromatin. DNA was counterstained with DAPI (blue). **d**, Line scans of the merged images of the cells in (**c**). The fluorescence intensity of H3K27me3, DNA (DAPI), and RING1B or BMI1 was scanned along the white arrows. **e**, Pearson’s correlation coefficients between H3K27me3 and CBX2/4/6/7/8/RING1B/BMI1. Data are presented as means ± SDs. Representative *p* values are for student *t*-test between each sample and CBX7 (****, *p*<0.0001; ns, not significant). **f**, Domain structures of the core subunits of the PRC1 complex. In mouse CBX7, the Chromodomain, AT-hook (AT), and Polycomb box (CBox) are shown. In human RING1B, the Ring finger domain at the N-terminus appears to be interacting with the RING domain of BMI1. RING-finger and WD40-associated ubiquitin-like (RAWUL) domain at the C-terminus, which interact with the CBox domain (represented by the dotted line). In human BMI1, the Low-Complexity Domain (LCD) is shown at the C-terminus. fPHC1 is the truncated version of *Drosophila* Ph-p (aa 1289-1577) fused with the HD1 domain (aa 23-52) of human PHC2 isoform B at the N-terminus. **g**, A model of the PRC1 complex based on the published information ^38^ and the gel-filtration data in (**h**). The SAM domain forms limited polymers of 4-6 units (6 are shown). **h**, Gel-filtration chromatograms of PRC1 complex (black) with an excess of CBX7/RING1B subcomplex. The protein standards are represented by the thin gray lines. **i,** SDS-PAGE analysis of the PRC1 complex in (**h**). **j**, A schematic showing the assembly of an Alexa Fluor 594-labeled H3K27me3 NA. **k**, Electron microscopy image of negatively stained H3K27me3 12xNA. **l**, Phase separation assay of PRC1 (5 μM) complex with reconstituted H3K27me3 12xNA (100 nM). **m**, A phase diagram of PRC1 complex with H3K27me3 12xNA. The corresponding images are shown in Extended Data Fig. 1c. Red means LLPS. The height of each bar corresponds to the percent area of the image occupied by droplets. PRC1 complex concentrations are in micromolar, and H3K27me3 12xNA concentrations are in nanomolar. **n**, Phase separation assay of PRC1 complex with reconstituted native and H3K27me3 12xNA. CBX7(F11A)-PRC1 or PRC1-SAM (L1547R/L1565R) complex was mixed with H3K27me3 12xNA. All complexes are at 5 μM.

We next studied interactions between CBX7-PRC1 and H3K27me3 chromatin *in vitro*. We obtained a recombinant CBX7-PRC1 complex containing CBX7, RING1B, MBP-tagged BMI1, and fPHC (Fig. 1f,g), in which fPHC1 contains a human homology domain 1 (HD1) to interact with human BMI1 at the N-terminus, and a truncated version of *Drosophila* Ph-p (termed Mini Ph-p) at the C-terminus. Mini Ph-p exists as polymers of 4-6 units via its SAM domain^8^. These four subunits co-elute as a stoichiometric complex before CBX7/RING1B on size-exclusion chromatography (Fig. 1h,i). Therefore, the potential reader valence in the CBX7-PRC1 complex is at least four or six, which is derived from the SAM domain of fPHC1. Human PHC1 is of higher oligomerization and it is challenging to obtain recombinant CBX7-PRC1 containing human PHC1.

To investigate the multivalent H3K27me3-reader interaction, we reconstituted nucleosome arrays (NA), referred to as 12xNA, composed of recombinantly purified and fluorophore-labeled histone octamers and a defined DNA template containing 12 repeats of Widom’s 601 nucleosome positioning sequence. For H3K27me3-marked NA, we adopted the methyl-lysine analog strategy^9^ to obtain structurally and biochemically similar analogs of H3K27me3 (Fig. 1j,k). When H3K27me3 12xNA was mixed with CBX7-PRC1, numerous puncta formed (Fig. 1l,m and Extended Data Fig. 1b).

We tested whether the puncta were resulted from LLPS driven by multivalent H3K27me3-reader interactions. Firstly, PRC1 formed puncta with H3K27me3 12xNA across a range of concentrations (Fig. 1n and Extended Data Fig. 1c). Secondly, we found that puncta fused when they contacted one another, then relaxed toward larger spherical ones (Extended Data Fig. 1d). Thirdly, we used the FRAP (fluorescence recovery after photobleaching) assay to show that PRC1 exchanged inside the puncta (Extended Data Fig. 1e,f), a property consistent with LLPS. We further investigated the role of multivalency by analyzing punctum formation using PRC1 with a mutated CBX7 (CBX7(F11A)), which cannot bind H3K27me3(ref. ^10^), and a double-mutated PHC1 (fPHC1(L1547R/L1565R)), which cannot polymerize^11,12^, to form CBX7(F11A)-PRC1 and PRC1-fPHC1(L1547R/L1565R), respectively (Extended Data Fig. 1g). Neither mutant PRC1 produced substantial puncta, if any, with H3K27me3 12xNA at the same reader concentrations (Fig. 1n). These results indicated that a critical multivalent H3K27me3-reader interaction drove LLPS.

When PRC1 compacts and stabilizes facultative heterochromatin *in vivo*, the DNA-contacted component of the chromatin is normally less accessible to external factors. PRC1 and H3K27me3 12xNA condensates were shown to be resistant to DNase I digestion (Extended Data Fig. 1h), suggesting that these *in vitro* phase-separated condensates resemble facultative heterochromatin *in cellulo* as far as DNA accessibility is concerned.

We previously demonstrated that constitutive heterochromatin is regulated, in part, via LLPS mediated by multivalent H3K9me3 and heterochromatin protein 1 (HP1)-containing complexes^1^. A recent study found that acetylated and non-acetylated chromatin form two distinct phases in the presence of poly(bromo)^3^. To better understand compartmentalization of chromatin within the nucleus, we focused on endogenous localizations of H3K27me3, H3K9me3, and their readers. Consistent with literature^2,13^, we found that H3K27me3-positive and H3K9me3-positive condensates in NIH-3T3 cells were mutually exclusive, and maintained their own compartments independently (Fig. 2a,b). CBX7 and HP1β strongly colocalize with H3K27me3 and H3K9me3, respectively (Figs. 1a, 2c-e). We next explored the cellular role of LLPS-mediated compartmentalization driven by multivalent PTM-reader interactions. Overexpression of CBX7(F11A) largely diminished H3K27me3-marked facultative heterochromatin, but had little effects on H3K9me3-marked constitutive heterochromatin; overexpression of HP1β(V23M), which has reduced H3K9me3-binding capacity, reduced H3K9me3-marked constitutive heterochromatin but had little effect on H3K27me3-marked facultative heterochromatin (Extended Data Fig. 2a,b). These results indicate that multivalent interactions of PTM-reader pairs are required for differential heterochromatic compartmentalization in cells.

**Fig. 2:**
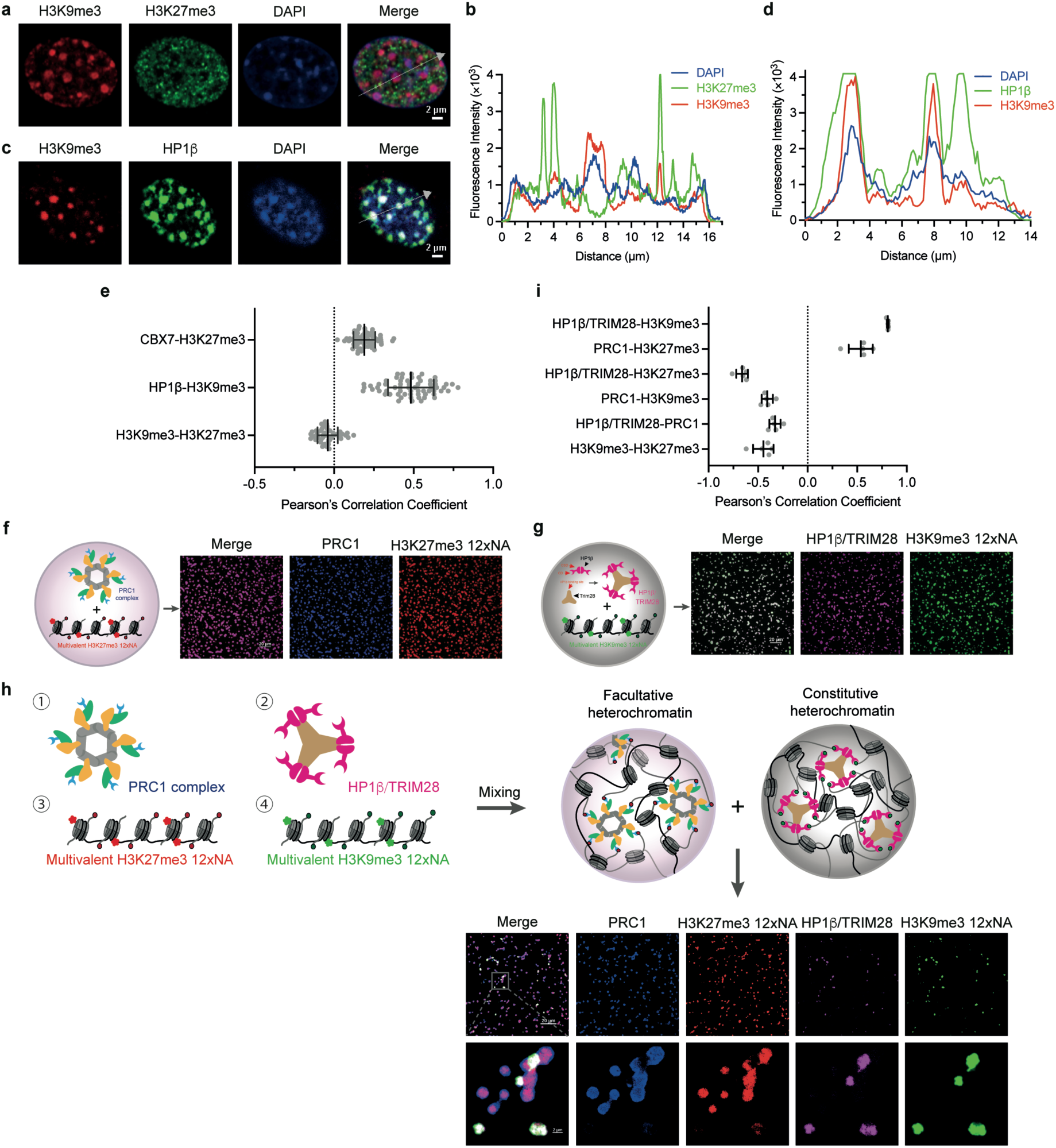
H3K27me3 and H3K9me3 form immiscible puncta *in cellulo* and *in vitro*. **a**, Immunofluorescence images of NIH-3T3 cells show that H3K9me3 (red) has no colocalization with H3K27me3 (green) in the nucleus. DNA was counterstained with DAPI (blue). **b**, Line scans of the merged image of the cell in (**a**). The fluorescence intensity of H3K9me3, H3K27me3, and DNA (DAPI) was scanned along the white arrow. **c**, Immunofluorescence images of NIH-3T3 cells show that H3K9me3 (red) colocalizes with HP1β (green) in the nucleus. DNA was counterstained with DAPI (blue). **d**, Line scans of the merged image of the cell in (**c**). The fluorescence intensity of H3K9me3, HP1β, and DNA (DAPI) was scanned along the white arrow. **e**, Pearson’s correlation coefficients between H3K9me3 and H3K27me3 (n=49); HP1β and H3K9me3 (n=64); CBX7 and H3K27me3 (n=64). All of the data are presented as means ± SDs. **f**, Left: a model showing the multivalent H3K27me3-PRC1 interactions which drive LLPS *in vitro*. Right: fluorescence images of *in vitro* H3K27me3-PRC1 condensates. PRC1 complex was partially (5%) labeled with Alexa Fluor 405 (blue), and H2B was totally labeled with Alexa Fluor 594 (red) in H3K27me3 12xNA. PRC1 concentration, 5 μM; 12xNA concentration, 500 nM. **g**, Left: a model showing the multivalent H3K9me3-HP1β/TRIM28 interactions which drive LLPS *in vitro*. Right: fluorescence images of *in vitro* H3K9me3-TRIM28 condensates. HP1β/TRIM28 was partially (5%) labeled with Alexa Fluor 647 (purple), and H2B was totally labeled with Alexa Fluor 488 (green) in H3K9me3 12xNA. HP1β/TRIM28 concentration, 10 μM; H3K9me3 12xNA concentration, 2 μM. **h**, The top row shows a schematic summary of droplet formation in (**f**) and (**g**). The images in the bottom row show that H3K27me3-PRC1 droplets are immiscible with H3K9me3-HP1β/TRIM28 droplets *in vitro*. The area in the gray box (top) is zoomed-in at the bottom (scale bar = 2 μm). PRC1 concentration, 5 μM; H3K27me3 12xNA concentration, 500 nM; HP1β/TRIM28 concentration, 1.25 μM; H3K9me3 12xNA concentration, 250 nM. **i**, Pearson’s correlation coefficients of PRC1, H3K27me3, HP1β/TRIM28, and H3K9me3 with each other. The corresponding values are the results of different protein concentrations. Representative fluorescence images are shown in (**h**) and Extended Data Fig. 2c. All of the data are presented as means ± SDs.

PRC1 and HP1β/TRIM28 can phase separate into liquid droplets with H3K27me3 12xNA and H3K9me3 12xNA, respectively (Fig. 2f,g)^1^. We next tested the condensation behaviors with H3K27me3 12xNA, H3K9me3 12xNA, and their readers together. When the four components were sequentially added to test tubes, we observed a three-phase system with H3K27me3-PRC1 condensates coexisting immiscibly with H3K9me3-HP1β/TRIM28 condensates, surrounded by the dilute buffer phase (Fig. 2h,i and Extended Data Fig. 2c). This condensate organization is consistent with what we observed in NIH-3T3 nucleoli (Figure 2e). FRAP experiments showed that both PRC1 and HP1β/TRIM28 signals exhibit partial liquidity within their condensates (Extended Data Fig. 2d,e). However, omitting H3K27me3 12xNA or H3K9me3 12xNA, the remaining three components co-condensate (Extended Data Fig. 2f-i). These results hint that dysregulation of epigenetic factors could affect chromatin compartmentalization.

To assess the molecular mechanism underlying the aforementioned immiscibility, we employed two strategies. First, we tempted to engineer model multivalent chromodomain systems. Yeast SmF, a 14-meric protein^14^ was previously used to establish multivalent scaffolds that can robustly drive LLPS^15^. Herein, we fused SmF with the chromodomain of CBX7 or HP1β to form 14-meric CD systems, named SmF-CD(CBX7) or SmF-CD(HP1β) (Fig. 3a). SmF-CD(CBX7) and SmF-CD(HP1β) formed condensates with H3K27me3 12xNA and H3K9me3 12xNA, respectively (Extended Data Fig. 3a). We next mixed these four components together and observed a three-phase system with H3K27me3/SmF-CD(CBX7) condensates coexisting immiscibly with H3K9me3/SmF-CD(HP1β) condensates (Fig. 3b). Subsequently, we swapped the CDs of HP1β and CBX7 and assembled the chimeric proteins into PRC1 and HP1β/TRIM28 complexes, called PRC1^CD(HP1β)^ and HP1β/TRIM28^CD(CBX7)^, respectively (Fig. 3c). We found that HP1β/TRIM28^CD(CBX7)^ and PRC1^CD(HP1β)^ can phase separate into condensates with H3K27me3 12xNA and H3K9me3 12xNA, respectively (Fig. 3d,e). When these four components were mixed together, H3K27me3-HP1β/TRIM28^CD(CBX^^7)^ condensates coexisted with H3K9me3-PRC1^CD(HP1β)^ condensates but the two were immiscible (Fig. 3f). These data indicated that the PTM-reader pairs dictate the specificity involving constitutive heterochromatin-like condensates and facultative heterochromatin-like condensates.

**Fig. 3:**
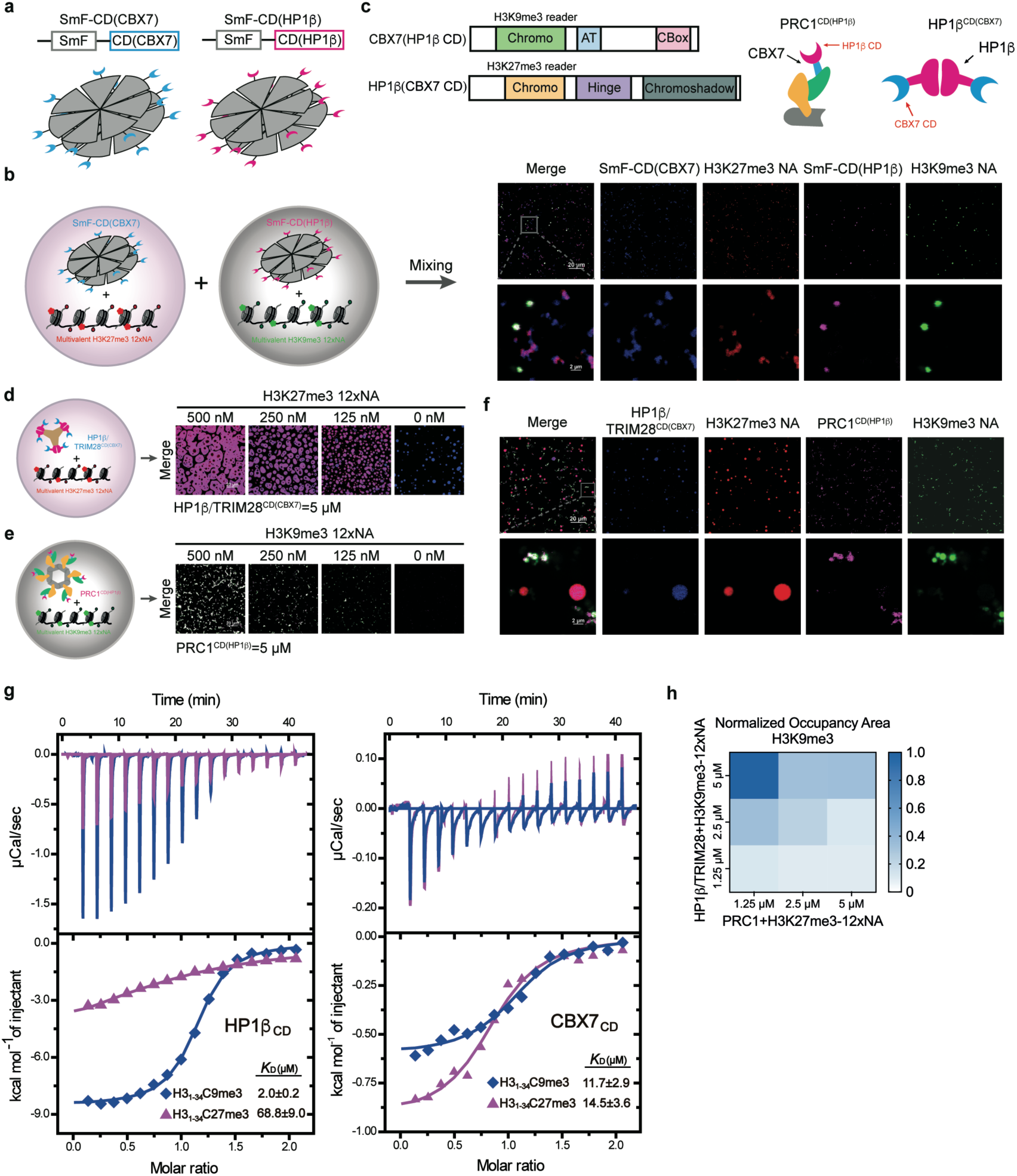
Mechanism of formation of immiscible compartments. **a**, Multimerization of chromodomains by fusion with a 14-meric protein, yeast SmF. CBX7 CD or HP1β CD is fused to the C-terminus of SmF. **b**, Left: a schematic summary of H3K27me3-SmF-CD(CBX7) droplets mixed with H3K9me3-SmF-CD(HP1β) droplets. Right: immiscible droplets *in vitro*. SmF-CD(CBX7) was partially (5%) labeled with Alexa Fluor 405 (blue), and H2B was totally labeled with Alexa Fluor 594 (red) in H3K27me3 12xNA. SmF-CD(HP1β) was partially (5%) labeled with Alexa Fluor 647 (purple), and H2B was totally labeled with Alexa Fluor 488 (green) in H3K9me3 12xNA. The region in the gray box is zoomed-in at the bottom. SmF-CD(CBX7) or SmF-CD(HP1β), 10 μM; H3K27me3 or H3K9me3 12xNA, 1 μM. **c**, Schematic of the chromodomain exchange of CBX7 and HP1β. Left: Domain structures of the exchanged proteins. CBX7(HP1β CD) is CBX7 with the chromodomain of HP1β and HP1β (CBX7 CD) is HP1β with the chromodomain of CBX7. Right: Models of the resulting exchanged complexes (PRC1^CD(HP1β)^ complex and HP1β^CD(CBX7)^ complex). **d**, Left: A model of the multivalent H3K27me3-HP1β/TRIM28^CD(CBX7)^ interactions. Right: Fluorescence images showing LLPS of HP1β/TRIM28 ^CD(CBX7)^ and H3K27me3 12xNA *in vitro*. HP1β/TRIM28^CD(CBX7)^ was partially (5%) labeled with Alexa Fluor 405 (blue), and H2B was totally labeled with Alexa Fluor 594 (red) in H3K27me3 12xNA. **e**, Left: A model of the multivalent H3K9me3-PRC1^CD(HP1β)^ interactions. Right: Fluorescence images showing LLPS of PRC1^CD(HP1β)^ and H3K9me3 12xNA *in vitro*. PRC1^CD(HP1β)^ was partially (5%) labeled with Alexa Fluor 647 (purple), and H2B was totally labeled with Alexa Fluor 488 (green) in H3K9me3 12xNA. **f**, Fluorescence images of H3K27me3-HP1β/TRIM28^CD(CBX7)^ droplets mixed with H3K9me3-PRC1^CD(HP1β)^ droplets. The area in the gray box is zoomed-in at the bottom. HP1β/TRIM28^CD(CBX7)^, 10 μM; H3K27me3 12xNA, 2 μM; PRC1^CD(HP1β)^, 5 μM; H3K9me3 12xNA, 500 nM. **g**, ITC curves of H3C9me3 and H3C27me3 peptides titrated into CD of HP1β (left) and CBX7. **h**, A heat map showing the normalized area occupied by H3K9me3 signal in the H3K9me3-HP1β/TRIM28 and H3K27me3-PRC1 mixing assays. HP1β/TRIM28 concentrations are labeled on the left, and H3K9me3 12xNA concentrations are one-fifth of HP1β/TRIM28. PRC1 concentrations are labeled at the bottom, and H3K27me3 12xNA concentrations are one-tenth of PRC1. The corresponding images are shown in Extended Data Fig. 3d. The color depth of each block corresponds to the percent area of the image occupied by droplets.

To gain more insights into the mechanism underlying condensate immiscibility, we measured the affinities of CBX7 CD and HP1β CD for the histone peptides in the reconstituted NAs (Extended Data Fig. 3b). We found that HP1β CD prefers H3K9me3 to H3K27me3 while CBX7 CD displays similar affinities towards H3K9me3 and H3K27me3 (Fig. 3g and Extended Data Fig. 3c). Our data suggest that the high-affinity interaction of HP1β and H3K9me3 prioritizes their binding to condensate, then CBX7 formed condensates with H3K27me3. From these results, we conclude that multivalent PTM-reader interactions, and their differences in binding preferences, mediate condensate immiscibility, which likely underlies the formation of co-existing chromatin compartments in nuclei.

When the H3K27me3-PRC1 system was mixed with the H3K9me3-HP1β/TRIM28 system *in vitro*, we unexpectedly found that the latter condensates were compromised in the presence of excessive amount of the former (Fig. 3h and Extended Data Fig. 3d). To test whether similar behaviors happen in cells, we transiently transfected CBX7 into NIH-3T3 cells and monitored the distribution of H3K9me3 and H3K27me3. CBX7 overexpression resulted in fewer chromocenters based on H3K9me3 staining but did not saliently affect the H3K27me3 compartment (Fig. 4a-e). Stable expression of mCherry-CBX7 also showed the same results (Extended Data Fig. 4a). The levels of other PRC1 subunits also increased along with CBX7 overexpression (Fig. 4f), indicating that the whole PRC1 is upregulated. Western blotting confirmed that the level of endogenous H3K9me3 was anti-correlated with that of CBX7 in stable cell lines, under conditions of high expression of CBX7 (Fig. 4f). Perplexingly, when the expression level of CBX7 was slightly increased, it seemed to have a promoting effect on H3K9me3 (Fig. 4f). This phenomenon warrants further studies.

**Fig. 4:**
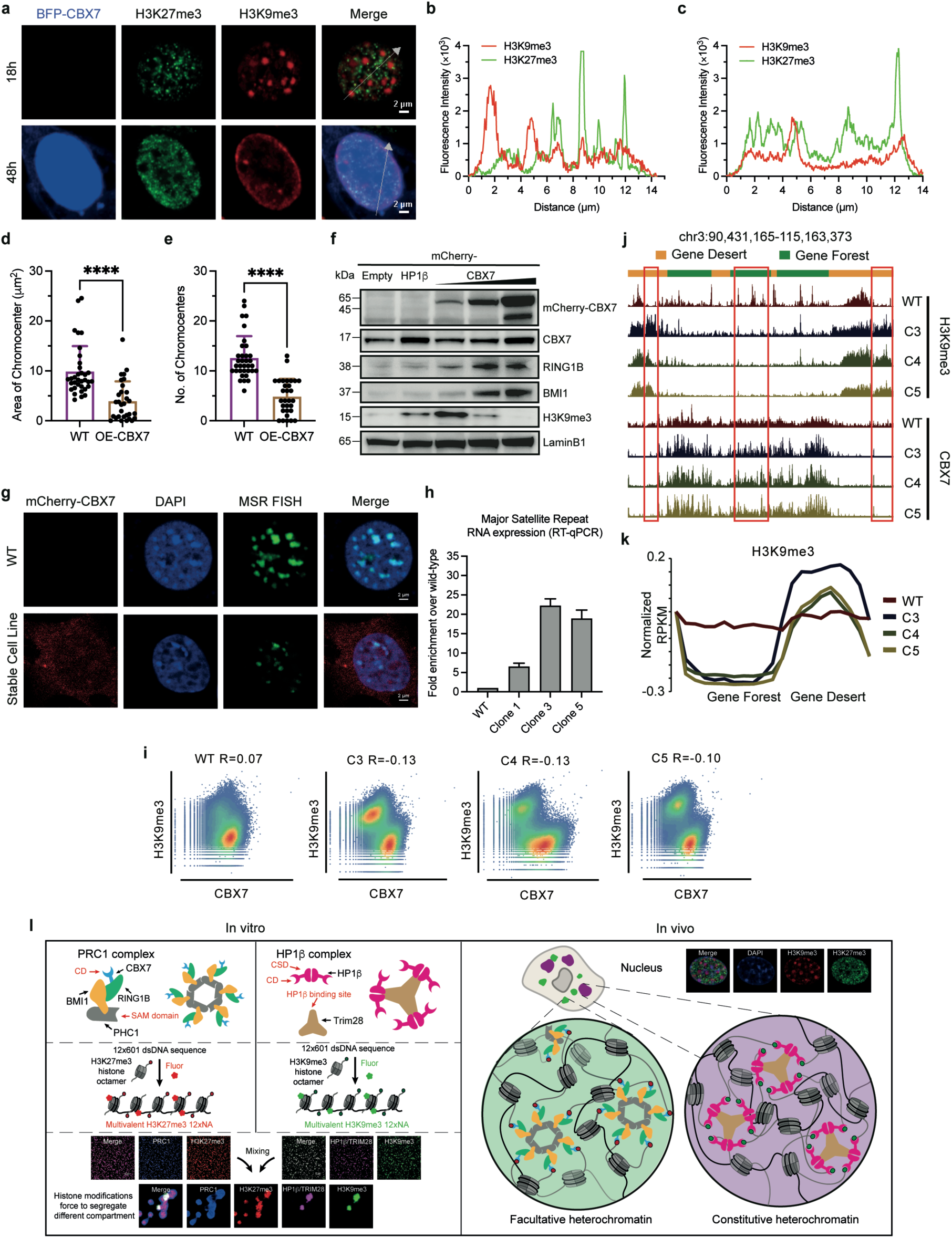
PRC1 over-expression compromises constitutive heterochromatin. **a**, CBX7 over-expression affects the distribution of H3K9me3 in NIH-3T3 cells. Immunostaining data show that the transient expression of BFP-CBX7 for 48 hours dissolves H3K9me3 (red) compared with transient expression for 18 hours. **b**,**c**, Line scans of the images of cells overexpressing BFP-CBX7 at 18 and 48 hours. Scanning was carried out along the white arrows in the merged images in (**a**). **d**,**e**, Area and number of chromocenters in native (WT) 3T3 cells compared with cells transiently expressing CBX7 (OE-CBX7) (n=34/30 cells). Data are presented as means+SDs and *p* values are for student *t*-test (****, *p*<0.0001). **f**, Western blotting analysis of NIH-3T3 cell lines with stable expression of mCherry-Empty, mCherry-HP1β, and mCherry-CBX7 in Extended Data Fig. 4a. LaminB1 loading was used as a control. **g**, Fluorescence images of major satellite repeats (MSR) (green) by DNA FISH in native NIH-3T3 cells (WT) and NIH-3T3 cells with stable expression of mCherry-CBX7. DNA is labeled by DAPI (blue). **h**, Reverse transcription-quantitative PCR (RT-qPCR) analysis of total RNA isolated from unsynchronized wild-type NIH-3T3 cells and three monoclonal NIH-3T3 cell lines overexpressing CBX7 to detect expression from the major satellite repeats with MSR-specific primers. The amplified signals were normalized to β-actin and are plotted in the histogram. Data are presented as means+SDs of at least two independent experiments, and *p* values are for student *t*-test (*, *p*<0.0332; ***, *p*<0.0002; ****, *p*<0.0001). **i**, Scatter plots showing the co-existence of CBX7 and H3K9me3 in WT cells and anti-correlation of CBX7 and H3K9me3 in three mCherry-CBX7 clones. The Pearson’s correlation coefficient is shown at the top. **j**, UCSC browser view showing the gain of H3K9me3 in the gene desert regions (yellow) and loss of H3K9me3 in the gene forest regions (green) upon CBX7 binding. Examples are enclosed in the red boxes. **k**, Average H3K9me3 enrichment (Z-score normalized RPKM) in gene desert regions and gene forest regions in wild-type cells and mCherry-CBX7 clones. **l**, Model showing how the formation of immiscible phase-separated condensates underlies chromatin compartmentalization.

To examine the reason for and the consequence of chromocenter dissolution, we visualized the positioning of major satellite repeats (MSR) by DNA fluorescence *in situ* hybridization (FISH) with fluorescence-tagged oligonucleotide probes that specifically target the consensus sequences of MSR. The abundance of heterochromatin clusters was reduced in mCherry-CBX7 NIH-3T3 cells, indicating a global dissolution of H3K9me3-marked chromocenter (Fig. 4g). Consistently, we analyzed transcription of MSR by RT-qPCR and found that there was significant de-repression of MSR in mCherry-CBX7 overexpression cells (Fig. 4h).

The redistribution of H3K9me3 might be caused by the dissolution of local H3K9me3/HP1 condensates after binding of CBX7-PRC1, as is shown by the antagonistic relationship between CBX7-PRC1 and H3K9me3 in mCherry-CBX7 3T3 cells (Fig. 4i). Therefore, we performed ChIP-seq (and CUT&Tag) and Hi-C experiments^16–18^ on WT and mCherry-CBX7 NIH-3T3 cells. We did not observe salient changes in H3K27me3, H2AK119Ub, and H3K4me3 ChIP patterns (Extended Data Fig. 4b). However, H3K9me3 tended to increase in gene desert regions and decrease in gene dense regions, where CBX7 predominantly binds (Fig. 4j,k and Extended Data Fig. 4c). We also observed disruption of chromatin higher order structure and transversion of A/B compartments in the cell lines (Extended Data Fig. 4d,e). To further determine the anti-correlation between H3K27me3-compartment and H3K9me3-compartment, we knocked out CBX7 via CRISPR/Cas9 in NIH-3T3 cells. CBX7 KO cells have more H3K9me3 compartments than wild-type cells (Extended Data Fig. 5a,b). These results together shed light on the roles of histone modifications and their regulatory complexes in the formation of constitutive and facultative heterochromatin: histone modifications act as anchors that mark where and which type of heterochromatin is to be constructed, while LLPS of these marks with their regulatory complexes controls the balance between constitutive and facultative heterochromatin.

Several studies have shown that overexpression of CBX7 and loss of H3K9me3 are related to the emergence of AML^19–21^, so we analyzed the transcriptome of mCherry-CBX7 3T3 cell lines. We detected up-regulation of leukemia-related genes^22–25^, and down-regulation of genes related to nucleosome assembly and genomic stability (Extended Data Fig. 6a-c). We also detected AML-associated genes that lost deposition of H3K9me3, but not other histone marks including H3K27me3 and H2AK119Ub (Extended Data Figs. 5c and 7a).

Subsequently, we wondered whether AML cells share a common phenotype of H3K9me3 compartment loss. Hematopoietic stem cells (HSCs) in mammals are positive with CD34 antigen (CD34^+^)^26^. CBX7 overexpression in HSCs enhances self-renewal and induces leukemia. This effect is dependent on integration of CBX7 into PRC1 and requires H3K27me3 binding^19^. We analyzed the distribution of H3K9me3 and H3K27me3 in two AML cell lines (THP-1 and OCI-AML3) and human CD34^+^ cord blood cells (Extended Data Fig. 7b). Strikingly, high CBX7 levels in THP-1 and OCI-AML3 were associated with a reduction of the size and abundance of chromocenters based on H3K9me3 staining (Extended Data Fig. 7c).

A small molecule MS37452 has been reported to bind CD of CBX7 and prevent its binding with H3K27me3 (refs. ^20,27^). We found that MS37452 reduced LLPS of CBX7-PRC1 and H3K27me3 12xNA *in vitro* (Extended Data Fig. 7d). Interestingly, the H3K9me3 compartment was partially reconstructed upon treatment with MS37452 (Extended Data Fig. 7e). We found that MS37452 induced a dramatic increase in the area and number of chromocenters in THP-1 and OCI-AML3 cells (Extended Data Fig. 7f-j). Then, we assessed to what extent inhibiting CBX7-PRC1 phase separation would affect leukemic cell growth. While HEK293 cells were insensitive to treatment with MS37452 up to 100 μM (Extended Data Fig. 7k), treatment with MS37452 resulted in compromise in proliferation to both THP-1 and OCI-AML3 cells in a dose-dependent manner (Extended Data Fig. 7l). These results suggest that restoring constitutive heterochromatin by inhibiting CBX7-PRC1 and H3K27me3 LLPS can reduce survival of AML cells.

We have previously demonstrated that LLPS driven by multivalent interactions between H3K9me3 and its chromodomain plays a significant role in the formation of constitutive heterochromatin^1^. Building upon our earlier work, this study reveals robust LLPS resulting from the interaction between H3K27me3 nucleosome arrays and its multivalent reader complex, CBX7-PRC1 (Fig. 1n). In addition, our data demonstrate that LLPS driven by multivalent H3K9me3-chromodomain and multivalent H3K27me3-CD interactions contributes to the formation of constitutive heterochromatin and facultative heterochromatin, respectively. Interestingly, the condensates derived from H3K9me3-CD coexist with those formed by H3K27me3-CD *in vitro* (Fig. 4l). This immiscibility of condensates is reminiscent of, and likely underlies, the segregation of constitutive heterochromatin and facultative heterochromatin in the nuclei.

Constitutive heterochromatin and facultative heterochromatin are regulated by H3K9me3 and H3K27me3, in conjunction with their corresponding chromodomain-containing reader complexes. Notably, the amino acid sequences of the H3K9 motif, ARKST, and that of the H3K27 motif, ARKSA, are highly similar. Additionally, CBX7 CD can bind both H3K9me3 and H3K27me3 and HP1 CD shows a preference for H3K9me3. Considering these two facts, the immiscibility of the two kinds of condensates is remarkable and serves as a clear manifestation of the high cooperativity specifically enabled by phase separation.

The cross-reactivity observed between H3K9me3-chromodomain and H3K27me3-CD pairs suggests a possibility of interconversion between these two heterochromatic compartments, both in normal and abnormal scenarios. Supporting this idea, we observed that overexpression of CBX7-PRC1 led to the reduction of constitutive heterochromatin compartments in NIH-3T3 cells. Numerous studies have reported data consistent with the interconversion of these two types of heterochromatic compartments during development across various species. For example, in the pericentromeric heterochromatin domain of embryonic stem cells, H3K9me3 and H3K27me3 exhibit mutual antagonism. ChIP-seq^28^ and staining data^29,30^ have shown that deficiency of H3K9me3 results in chromatin reorganization, wherein H3K27me3 undergoes a localized transition into regions otherwise enriched with H3K9me3 in wild-type cells.

Regarding abnormal interconversion between constitutive heterochromatin and facultative heterochromatin, it is well-known that CBX7 tends to be overexpressed in AML cells^20^. Our study provides valuable epigenetic mechanistic insights into the pathogenesis of AML. MS37452 effectively weakens the phase separation of PRC1 with H3K27me3 12xNA, presenting an opportunity for the re-establishment of H3K9me3 condensates. CBX7 has been reported to modulate lifespan and delay cellular senescence when ectopically expressed^31^, it will be interesting to determine whether the reconstruction of heterochromatin causes senescence-associated heterochromatic foci that inhibit cancer proliferation.

Essentially the high degree of cooperativity associated with switch-like phase separation enables a clear distinction between “seemingly promiscuous” biochemical systems, i.e. systems with subtle differences in microscopic interactions. Our study also provides potential epigenetic mechanistic insights into tumor development and suggests the possibility of regulating phase separation for cancer therapy.

## Methods

### Cell lines

NIH-3T3 cells were maintained in DMEM (SH30022.01, Hyclone) containing 5% FBS and 100 units/mL penicillin-100 μg/mL streptomycin solution (SV30010, HyClone).

THP-1 and OCI-AML3 cells were maintained in RPMI 1640 Medium (61870036, Thermo Fisher) containing 10% FBS (10099141C, GIBCO), 100 units/mL penicillin-100 μg/mL streptomycin solution.

### Cloning

The pRSFDuet1-His6-HP1β plasmid and pETDuet1-His6-MBP-TRIM28 (1-501) have been described previously ^1^. The sequence of HP1β^CD(CBX^^7)^ in pRSFDuet1-His6 was the substitution of *H. sapiens* HP1β Chromo Domain (18-72) with *M. musculus* CBX7 Chromo Domain (8-69). We cloned *H. sapiens* HP1β CD, *M. musculus* CBX7, CBX7F11A, or CBX7^CD(HP1β)^ (substitution of *M. musculus* CBX7 Chromo Domain with *H. sapiens* HP1β Chromo Domain) with His6-MBP-tag followed by an *N. tabacum* etch virus protease ^32^ recognition site into the MCS1 of pRSFDuet1 expression vector. For CBX7/RING1B subcomplexes (wild-type, truncation, and mutation), *H. sapiens* RING1B with a His6-tag followed by a TEV protease recognition site was cloned into MCS2 of this vector. We cloned *H. sapiens* BMI1 with His6-MBP-tag followed by a TEV protease recognition site into the MCS1 of pRSFDuet1 expression vector, *Drosophila* Ph-p (1289-1577) or Ph-p (L1547R/L1565R) mutation fused with *H. sapiens* PHC2B HD1 domain (23-52) at the N terminus was cloned with a His6-tag followed by a TEV protease recognition site, into MCS2 of this vector.

The *S. cerevisiae* SmF variants with His6-MBP-SUMO-tag followed by a SUMO protease 1 (Ulp1) recognition site were constructed into the pRSFDuet1 together with CBX7 CD or HP1β CD. CBX7 CD (1-73) with His10-SUMO-tag followed by Ulp1 recognition site was constructed into the pSUMOH10 expression vector.

Plasmids for the expression of wild-type H2A, H2B, and H3K27C/C110A histone proteins from *X. laevis* were generous gifts from Dr. Guohong Li. We engineered plasmid for the expression of *X. laevis* H2BT116C with EasyMut gene multipoint mutation kit (CL307-01, Biomed). Plasmids for the expression of wild-type *X. laevis* H4 histone proteins were a generous gift from Dr. Leiming Xie.

### Protein expression and purification

All proteins were expressed in *Escherichia coli* BL21 (DE3) cells. HP1β/TRIM28 complex purification has been described previously ^1^, same as the purification of HP1β CD and HP1β /TRIM28^CD(CBX^^7)^ proteins. The CBX7/RING1B (wild-type, mutation, or HP1β CD substituted CBX7 CD) subcomplex was co-expressed. Cells were grown to OD 0.8 at 37°C in LB media with 30 μg ml^−1^ Kanamycin. *E. coli* cells expressing the protein/protein complexes were induced with 1 mM isopropyl-β-d-thiogalactopyranoside (IPTG) at 16°C overnight, spun down at 4,000 rpm, and resuspended in lysis buffer (40 mM Tris-HCl pH 8.0, 500 mM NaCl, 10% Glycerol, 1 mM PMSF, 1 mM DTT), and lysed with a high-pressure homogenizer before centrifugation. The supernatants were first purified using Ni-NTA agarose. The His6-tag or His6-MBP-tag proteins were further purified by HiTrap Q anion-exchange, then cleaved by incubating with TEV protease overnight at 4°C. The cleaved sequence and uncleaved proteins removed by passing through a Ni-NTA agarose, and CBX7/RING1B subcomplexes were further purified using gel-filtration chromatography (SD200), then finally stored in 20 mM HEPES pH 8.0, 200 mM NaCl, 5 % Glycerol and 2 mM DTT before assembling PRC1 complex.

For SmF-CD(HP1β) and SmF-CD(CBX7) proteins purification, cells were induced with 1 mM isopropyl-β-d-thiogalactopyranoside (IPTG) at 18°C overnight, spun down at 4,000 rpm, and resuspended in lysis buffer (40 mM Tris-HCl pH 8.0, 500 mM NaCl, 5 % Glycerol, 1 mM PMSF, 1 mM DTT), and lysed with a high-pressure homogenizer before centrifugation. The supernatants were purified using Ni-NTA agarose, HiTrap Q anion-exchange, then cleaved by incubating with Ulp1 protease overnight at 4°C. The cleaved sequence and uncleaved proteins removed by passing through a Ni affinity chromatography, and further purified using gel-filtration chromatography (SD200), then finally stored in 20 mM HEPES pH 8.0, 500 mM NaCl, 5 % Glycerol, and 2 mM DTT at -80°C after being flash-frozen in liquid nitrogen. For CBX7 CD protein expression for ITC experiments, cells were growth in TB media at 37°C and switched to 15°C until OD reached 1.8-2.0. Cells were induced by 1 mM IPTG for 16-20 hours, spun down at 4,000 rpm, collected and lysed in PBS buffer containing 500 mM NaCl and 5% glycerol using high-pressure homogenizer. The lysate was centrifuged at 12,000 rpm for 1 hour and the supernatants were first purified using Ni-NTA agarose. His10-SUMO tag was cleaved by incubating with Ulp1 protease overnight at 4°C. The protein was purified using HiTrap Q anion-exchange column, gel-filtration chromatography (SD75), and finally stored in 30 mM HEPES pH 7.4, 100 mM NaCl before use.

The MBP-BMI1/fPHC1 (wild-type or mutation) subcomplex was co-expressed in *E. coli* Rosetta (DE3) strain. Cells were cultured at 37°C to OD600 0.8 and induced at 16°C for an additional 20 hours with 1 mM IPTG. Cells were harvested and lysed in binding buffer (40 mM Tris-HCl pH 8.0, 500 mM NaCl, 10mM β-ME, 100μM ZnCl2, 10% Glycerol, 1 mM PMSF, 1 mM DTT) and lysed with a high-pressure homogenizer before centrifugation. The supernatants were first purified using Ni affinity chromatography. The His6-tag or His6-MBP-tag proteins were further purified by HiTrap Q anion-exchange, then purified using Superpose 6 column, and finally stored in 20 mM HEPES pH 8.0, 500 mM NaCl, 10 % Glycerol, and 2 mM DTT at -80°C after being flash-frozen in liquid nitrogen.

For PRC1 (wild-type or mutation) complexes, 5 M NaCl solution was added to 500 mM final concentration to CBX7/RING1B subcomplexes in storage buffer (20 mM HEPES pH 8.0, 200 mM NaCl, 5 % Glycerol, 2 mM DTT), and mixed with the equal molar MBP-BMI1/fPHC1 subcomplexes followed by incubation for 1 h on the ice, then PRC1 complexes were concentrated in centrifugal concentrators with a 3,000 dalton MWCO to less than 500 μL per liter bacterial expression. Complexes were further purified using gel-filtration chromatography (SD200), then finally stored in 20 mM HEPES pH 8.0, 500 mM NaCl, 10 % Glycerol, and 2 mM DTT at -80°C after being flash-frozen in liquid nitrogen.

Recombinant histones from *X. laevis* were expressed and purified from *E. coli* as previously described ^33^, with some modification. Briefly, washed inclusion bodies containing *E. coli*-expressed histone proteins were soaked with 1 mL DMSO for 30 min at room temperature, and then solubilized with Unfolding buffer (50 mM Tris-HCl pH 8.0, 100 mM NaCl, 6 M Guanidinium•HCl, 10mM EDTA, 10 mM DTT) overnight at room temperature. The pellet should eventually almost completely dissolve, and centrifuge for 60 min at 4°C and 12,000 rpm, the supernatant contains the unfolded proteins. Pooled and dialyzed unfolded soluble inclusion body proteins three times against 0.1% TFA (85173, Thermo), remove white debris by centrifugation for 60 min at 4°C and 12,000 rpm. Proteins in the supernatant were further purified by HiTrap Heparin HP column (GE Healthcare) with A Buffer (50 mM Tris-HCl pH 8.0, 8 M Urea, 1 mM EDTA) and B Buffer (50 mM Tris-HCl pH 8.0, 8 M Urea, 1 mM EDTA, 1 M NaCl), flash-frozen in liquid nitrogen, and finally stored at -20°C.

### Protein labeling

PRC1 proteins were labeled by incubating with a 1:1 molar ratio of Alexa Fluor 405, Alexa Fluor 488 carboxylic acid (succinimidyl ester) (Thermo) for 1 hour at room temperature in dark. To remove the free dye, the samples were centrifugated through desalting columns (89882, Thermo), and the labeled proteins were finally stored at -80°C. For experiments requiring labeled PRC1 proteins, 10% labeled PRC1 was mixed with the complexes before use. HP1 and SmF-tagged proteins were labeled using the same process.

Histone H2BT116C was labeled as described previously ^3^.

### Preparation of the methyl-lysine analog (MLA) of histone H3K9me3 and H3K27me3

We adopted the methyl-lysine analog (MLA) strategy to obtain a structurally and biochemically similar analog of H3K9me3 or H3K27me3, in which a tri-methyl-aminoethyl group was introduced to the thiol group of cysteine 9 or 27 in the context of H3K9C/C110A or H3K27C/C110A histone H3. Briefly, 10 mg lyophilized H3K9C/C110A or H3K27C/C110A H3 sample powder was solubilized in 980 μL solubilization buffer (1 M HEPES pH 7.8, 4 M guanidine hydrochloride, 20 mM D/L-methionine, 20 mM DTT) by incubation at 37°C for 1 h. Then, 100 mg bromocholine bromide (TCI) was added to the solution to initiate the chemical reaction. After incubation at 50°C for 2 h, 10 μL of 1 M DTT was added and incubated for another 2 h. Repeat this chemical reaction. After that, 50 μL β-ME was added to quench the unconsumed bromocholine bromide, and stored at -20°C before octamer reconstitution. The tri-methylated lysine 9 and 27 analogs on H3K9C/C110A and H3K27C/C110A proteins were confirmed by western blot assays using monoclonal anti-Histone H3K9me3 and H3K27me3 antibody (Abcam).

### Preparation of DNA template

Preparation of the 12 × 177 bp 601 DNA template followed the method described previously ^1^ using a plasmid containing a DNA template harboring 12 × 177 bp tandem repeats of the Widom 601 sequence ^34^. The sequence of the 177 bp DNA repeat is as follows, with the 601 DNA sequence: GAGCATCCGGATCCCCTGGAGAATCCCGGTGCCGAGGCCGCTCAATTGGTCGTAGACAGCTCTAGCACCGCTTAAACGCACGTACGCGCTGTCCCCCGCGTTTTAACCGCCAAGGGGATTACTCCCTAGTCTCCAGGCACGTGTCACATATATACATCCTGTTCCAGTGCCGGACCC

### Reconstitution of nucleosomal arrays

Histone octamers were reconstituted using the method of serial dialysis. Briefly, equal molar amounts of individual histones were combined and dialyzed against refolding buffer (20 mM Tris-HCl pH 8.0, 2 M NaCl, 1 mM EDTA, 5 mM β-ME) two times for 3 h at 4°C and one time overnight at 4°C. After that, centrifuged for 30 min at 4°C and 12,000 rpm, and the supernatant contains the octamers. Assembled octamers were further purified through a Superdex 200 10/300 GL column (GE Healthcare), flash-frozen in liquid nitrogen, and finally stored at -80°C.

Reconstitution of nucleosomal arrays was performed at 4°C using the salt dialysis method as previously reported with minor modifications. DNA templates harboring 12 × 177 bp (1290275.5 g•mol^-1^) tandem repeats of the Widom 601 sequence were mixed with histone octamers at molar ratios of 1.1:1 (12 × 601 DNA: octamers = 1.1:1), and dialyzed over 18 hours in 450 mL 1×TE Buffer B (1 mM Tris-HCl pH 8.0, 2 M NaCl, 1 mM EDTA), which was continuously diluted by slowly adding 1050 mL 1×TE Buffer A (1 mM Tris-HCl pH 8.0, 1 mM EDTA) to decrease the NaCl concentration from 2 M to 0.6 M. The reconstituted 12 × nucleosome chromatin was further dialyzed against HE buffer (10 mM HEPES pH 8.0, 0.1 mM EDTA) for 4 hours before use.

### Microscopy

Imaging was done with a NIKON A1 microscope equipped with a 100 × oil immersion objective. NIS-Elements AR Analysis was used to analyze the images.

### DNase I treatment

For the treated (before phase separation (PS)) samples, 0.5 U DNase I (Transgene, GD201) and DNase I buffer were mixed with H3K27me3 nucleosomal arrays at 37°C for 15 min, then mixed with PRC1 complex. For the treated (after PS) samples, PRC1 complex was mixed with H3K27me3 nucleosomal arrays, then 0.5 U DNase I and DNase I buffer were added for 15 min at 37°C. For the control samples, PRC1 complex was mixed with H3K27me3 nucleosomal arrays, then DNase I buffer was added for 15 min at 37°C. The final concentration of PRC1 complex was 5 μM and the final concentration of H3K27me3 12 × nucleosomal arrays was 0.5 μM.

### Metal shadowing and electron microscopy

Before grid preparation, all samples were co-dialyzed against the same buffer (20 mM HEPES-Na pH 7.4, 100 mM NaCl) overnight. Holey grids coated with continuous carbon film were glow discharged, and then 4 µl of the sample were loaded. The grids were first blotted using filter paper, then washed with water and uranyl formate, sequentially. Samples were observed using a 120kV FEI Tecnai Spirit Bio TWIN microscope operating at a magnification of ×49,000.

### Phase separation assays

Preparation of 384-well microscopy plates, including mPEGylation of silica and passivation of well with Bovine Serum Albumin (BSA) were previously described ^3^. For PRC1 and 12 × nucleosomal arrays (NA) phase separation, we first diluted the NA in Phase Separation Buffer 0 / 500 (20 mM HEPES pH 8.0, 0 mM / 500 mM NaCl), then added PRC1 complex, the final NaCl concentration was 50 mM. Gently mixed and added to the well of a PEGylated and BSA passivated 384-well microscopy plate. HP1β/TRIM28 and H3K9me3 12 × NA phase separation assay was the same way.

### Fluorescence recovery after photobleaching (FRAP) measurements

In vitro FRAP experiments were carried out with a NIKON A1 microscope equipped with a 100 × oil immersion objective. Droplets were bleached with laser pulse (3 repeats, 80% intensity, dwell time 4 s). Recovery from photobleaching was recorded for the indicated time.

### Cell transfection

For transient transfections, when the cells reached 70% confluence, plasmids were transfected into NIH-3T3 cells using Lipofectamine 3000 reagent (L3000015, Invitrogen). The final concentration of plasmids was 1 μg/mL. 48 h after transfection, cells were imaged using a confocal microscope or fixed with 4% paraformaldehyde for immunofluorescence staining.

To construct NIH-3T3 cell lines stably overexpressing CBX7 or HP1β as cell models for functional study and sequencing, genes were amplified by PCR and ligated with the lentivirus expression vector pLVX-Puro (0.5 μg/mL), which was co-transformed into 293T cells using 0.25% [v/v] PEI transfection reagent with another three vectors, pPRE (0.25 μg/mL), pPEV (87.5 ng/mL), and pVSV (125 ng/mL). Replace with fresh complete medium after 24 h, after another 24 h, the culture medium was collected as virus solution, filtered, and then added to NIH-3T3 cells. Stable cell lines were screened with puromycin and sorted different expression levels or monoclones by BD FACSAria SORP. Cells were seeded in a 4-chamber glass bottom dish (D35C4-20-1-N, Invitrogen) for confocal imaging or in a 60 mm × 15 mm dish (705001, NEST) for western blotting.

### Genome editing with CRISPR/Cas9

NIH-3T3 cells were transfected with (px458, addgene 48138) containing sgRNA sequence targeting *Cbx7*. After FACS of GFP positive 3T3 cells, single cells were seeded, colonies were manully picked after 2 weeks and subsequently screened by PCR. DNA of 3T3 cells was isolated with QuickExtract^TM^ DNA Extraction Solution 1.0 (QE09050, Lucigen). Presence of the homozygous deletion was confirmed by PCR using specific primers. To confirm loss of CBX7 protein, cells were lysed in RIPA buffer, and detected by western blotting.

Primers for sgRNA assembly:

px458-Cbx7-up FWD: CACCGTTCAAAAATGGAACGTCGG

px458-Cbx7-up REV: AAACCCGACGTTCCATTTTTGAAC

px458-Cbx7-down FWD: CACCGCCCGACGGTTGTTTACAAAG

px458-Cbx7-down REV: AAACCTTTGTAAACAACCGTCGGGC

### CD34+ cord blood isolation

Cord blood was obtained from healthy full-term pregnancies after informed consent in accordance with the Declaration of Helsinki at The Fifth Medical Center of the PLA General Hospital (Beijing, China). The protocols were approved by the institutional ethics review boards from the Institute of Hematology and Blood Diseases Hospital, Chinese Academy of Medical Science and Peking Union Medical College. Initially, cord blood volume was measured and then diluted 1:1 with PBS+2% FBS+200 units/mL penicillin-200 μg/mL streptomycin solution (SV30010, HyClone). Maximum 25 mL of diluted core blood was carefully layered on 15 mL of Lymphoprep in a 50 mL SepMate^TM^ tube and centrifuged for 15 min at room temperature, 800 g, without brakes. Middle layer containing mononuclear cells was harvested and diluted 1:1 with Dilution Buffer (PBS+2% FBS+0.5 mM EDTA+200 units/mL penicillin-200 μg/mL streptomycin solution) and then centrifuged for 8 min at 1,000 g. Cell pellets were collected and washed with Dilution Buffer for 8 min at 1,000 g. Cell pellets were collected and resuspend by 500 μL (for 80 mL cord blood) Dilution Buffer and then incubated by 10 μL CD34-APC antibody (343510, BioLegend) on ice for 30 min. Cell suspension was washed with Dilution Buffer and centrifuged for 5 min at 1,000 g. Cell pellets were collected and resuspend by 500 μL Dilution Buffer and then incubated by 10 μL anti-APC microbeads (130-090-855, Miltenyi) on ice for 30 min, and then cells were enriched by filtering them through a 40 μm nylon mesh into the MACS columns. Collection CD34+ cells by centrifuged for 5 min at 1,000 g, cell pellets were collected and resuspend by Dilution Buffer containing 1μg/mL DAPI solution (BN20923), and then CD34+ living cells were sorted by BD FACSAria SORP.

### Immunofluorescence staining

For CD34+ HSCs, THP-1, and OCI-AML-3 suspension cell lines, first treated the glass bottom with the cell-attachment agent (C1010, APPLYGEN) at 37°C for 30 min, then seeded cells at 37°C for another 30 min. Cells were fixed with 4% paraformaldehyde for 10 min, then washed with PBS. Cells were then permeabilized with Triton X-100 (P0096, Beyotime) for 15 min and incubated for 1 h with 5% BSA. Cells were incubated with primary antibodies overnight at 4°C, then washed with PBS and incubated for 1 h with secondary antibodies. Cells were then washed with PBS and stained with DAPI (5 mg/L) for 10 min. After washing cells with PBS, images were captured by a Nikon A1RMP confocal microscope with a 100 × oil objective.

### Western blotting

The precipitates were resuspended using PBS and SDS loading buffer. Proteins were fractionated by SDS-PAGE and transferred to a PVDF membrane. The membranes were incubated overnight with primary antibodies, then with the corresponding peroxide-labeled IgG (1:5000) for 1 h. Finally, chemiluminescence reagent (#34077, Thermo Scientific) was used to visualize the results.

### Major Satellite Repeat-Fluorescence In Situ Hybridization (MSR-FISH)

Cells were plated on the poly-lysine coated cell slides (TW20-PLL, Twbio) in the 12-well plate (712001, NEST) overnight, and discarded the medium, washed with PBS once. Cells were fixed with 4% paraformaldehyde for 10 min at room temperature, then washed with PBS for 5 min twice. Cell membrane and nucleus membrane were permeabilized by methanol for 5 min, then washed with PBS for 5 min once. After that, incubated cells using 100 μg/mL RNase A in PBS for 30 min at 37°C, then washed with PBS for 5 min twice at room temperature. Cells were then heated on a hot plate at 80°C for 10 min in preheated (80°C) 80% formamide (Sigma-Aldrich, diluted in water), Alex488-tagged probes (a generous gift from Dr. Ge Zhan) in hybrid buffer were also heated at 80°C for 10 min to open high-order structure. Probe mixture contains 200-250 nM oligo probes in hybrid buffer (50% formamide, 2 × SSC, 20% dextran sulfate). Discarded 80% formamide and added 200 μL probe mixture to the slides, then incubated at 80°C for 10 min to make sure high-order structure was opened. Taked out slides and cool down to room temperature for 10 min, after that washed with 2 × SSCT (2 × SSC, 0.1% Tween) at 50°C for 15 min twice, then washed with 2 × SSCT for 5 min once at room temperature. Cells were incubated with DAPI (0.5 mg/mL in PBS) for 10 min at room temperature, dropped Fluoromount-G (0100-20, SouthernBiotech) onto the glass slides, and putted the slides face-down onto it, stored in the dark before imaging.

### Reverse Transcription quantitative PCR (RT-qPCR)

Total RNA was extracted with RNAiso plus (9108, TAKARA), then incubated with 5×gDNA wiper Mix and reverse transcribed using HiScript III 1st Strand cDNA Synthesis Kit (+gDNA wiper) (R213-02, Vazyme). Primers for the specific amplification of repeat element (MSR) and housekeeping gene (GAPDH) underlined:

MajSat – F: 5’-TGGAATATGGCGAGAAAACTG

MajSat – R: 5’-AGGTCCTTCAGTGGGCATTT

GAPDH-F: 5’-GGAGCGAGATCCCTCCAAAAT

GAPDH-R: 5’-GGCTGTTGTCATACTTCTCATGG

### Cell proliferation assay

CCK8 assay was conducted in accordance with the Cell Counting Kit-8 instructions (C0039, Beyotime). In brief, cells were seeded at a density of 3,000 cells/well into 96-well plates (701001, NEST) and treated with DMSO or MS37452 (SML1405, Sigma-aldrich). After days 2, 4, and 6, 10% [v/v] of CCK8 solution was added to each well of the plate, and the well was incubated for at least 2 h. Absorbance at 450 nm was evaluated using Biotek ELx808IULALXH. For dose-response studies, MS37452-treated cells were normalized to the DMSO-treated cells, the EC50 was calculated using the “log[inhibitor] vs. the normalized response-variable slope” equation in GraphPad Prism 9.

### Isothermal titration calorimetry (ITC)

All calorimetry experiments of the HP1β and CBX7 CDs were performed at 25°C using a MicroCal PEAQ-ITC instrument (Malvern Panalytical). Briefly, the CD proteins and the synthetic premodified H3 peptides (SciLight Biotechnology, LLC) were dialyzed in the following buffer: HP1β CD (20 mM HEPES, 150 mM NaCl, 1 mM TCEP, pH 8.0), CBX7 CD (30 mM HEPES, 100 mM NaCl, pH 7.4).

Peptides (>95% purity) were quantified by weighing in large quantities and then aliquoted and freeze-dried for future use. Protein concentration was determined by absorbance spectroscopy at 280 nm. Titration was performed by injecting peptides into the HP1β CD sample at the molar ratio of 1:0.1. The same titration was performed by injecting peptides into the CBX7 CD sample at the molar ratio of 2:0.2. Acquired calorimetric titration curves were analyzed using Origin 7.0 (OriginLab) using the “One Set of Binding Sites” fitting model. The raw data are shown in Table S1.

### ChIP-seq of CBX7

CBX7 ChIP-seq for WT and CBX7-mCherry cell lines were performed as previously described with modifications ^35^. 30 μL Dynal protein A beads (Thermo) were used per IP and were washed 1 time with 1 × PBS containing 5 mg/mL BSA before use. Beads were resuspended in 1ml BSA containing PBS buffer and with the tube against the magnet, 3 μg/IP antibody was added to the tube and the antibody and beads were incubated for more than 20 min at 4°C with rotating. After incubation, the antibody-conjugated beads were washed 1 time with BSA containing PBS buffer before use.

Cultured cells were digested with trypsin, the medium was removed by centrifuge, and cells were resuspended in 1×PBS. Cells were then centrifuged and resuspended in 1×PBS containing 1 % formaldehyde (Sigma) for 10 min at room temperature for fixation. Glycine was then added to a final concentration of 0.14 M and incubated for 10 min at room temperature to quench fixation. Cells were pelleted and resuspended in 800 μL cell lysis buffer (50 mM HEPES, pH 7.9; 5 mM MgCl2; 0.2% Triton X-100; 20% glycerol; 300 mM NaCl and freshly added ROCHE cOmplete protease inhibitor cocktail at 0.01 tablet/mL), transferred to a new 1.5 ml tube and incubated 10 min at 4°C with rotating. After removal of the supernatant, cells were resuspended in 400 μL SDS lysis buffer (0.3% SDS, 50 mM Tris-HCl pH 8.0, 20 mM EDTA, and fresh protease inhibitors) and rotated at 4°C for 10 min. The lysate is sonicated using the program configured as power 10, cycle 12, 15 seconds sonication, and 45 seconds rest for every minute). The lysates were then clarified by centrifuge at 4 °C 1.5 min at 13,000 rpm. 2 times of superDB buffer (50 mM Tris-HCl pH 8.0, 1.5% Triton X-100, 225 mM NaCl, 0.15% sodium deoxycholate and freshly added ROCHE cOmplete protease inhibitor cocktail at 0.01 tablet/mL) were added to the lysates and tubes were incubated at 4°C for 30 min with rotation.

40 μL pre-washed dynal beads (3 times using SDS lysis buffer) were added to the chromatins and rotated for 2 h at 4°C to pre-clear chromatin. After removing supernatant from antibody-bound beads, the pre-cleared sonicated chromatin was added to the tubes and they were incubated overnight at 4°C with rotating. The beads were then washed with cold buffers: twice with 1 mL 0.1% SDS lysis buffer, twice with 1mL high salt buffer (50 mM HEPES, pH7.5; 1 mM of EDTA; 1% Triton X-100; 0.1% sodium deoxycholate; 0.1% SDS; 350 mM of NaCl), twice with 1 mL of LiCl wash buffer (10 mM Tris-HCl, pH 8.0; 1 mM of EDTA; 0.5% NP-40; 0.5% sodium deoxycholate; 250 mM LiCl), and once with 1 mL TE Buffer. 100 μL elution buffer (10 mM EDTA, 1% SDS, 50 mM NaHCO3) was then added to beads, and tubes were incubated for 4 h at 65°C with vigorous shaking on a thermomixer. The supernatants were transferred to new tubes, 6 μL of 5 M NaCl was added to the tubes, and tubes were incubated overnight at 65°C for reverse crosslinking. 5 μL RNaseA was then added to each sample and incubated at 37°C for 1 h. 5 μL of proteinase K was then added to each tube and tubes were incubated 1 hour at 65°C. Nucleotides were purified using 2.5 × DNA beads (Vazyme) and eluted with 25 μL water. 1 μL Rnase A was added to each PCR tube and tubes were incubated for 1 h at 37°C. DNA were purified using 2.5 × DNA beads (Vazyme) and eluted with 25 μL water.

Tru-seq library preparation was done using NEBNext® Ultra TM II DNA Library Prep Kit for Illumina® v1.0 with some following modifications. In the end repair step, 25 μL eluted DNA, 3.5 μL reaction buffer and 1.5 μL enzyme mix were mixed in PCR tubes and incubated in a thermocycler. The program of the thermocycler was set as 20°C 30 min, 65°C 30 min, and 4°C hold. 1.25 μL of 0.6 μM adaptor, 15 μL ligation master mix, and 0.5 μL of ligation enhancer were added to each tube and mixed and tubes were incubated in a thermocycler. The program of the thermocycler was set as 4°C 2 h, 20°C 30 min, and 4°C hold. 1.5 μL of USER enzyme was then added to each tube and tubes were incubated in a thermocycler. The program of the thermocycler was set as 37°C 15 min, 4°C hold. Post-ligation cleanup was performed using 1.4×DNA beads (Vazyme) and eluted with 25 μL water. 10 μL adaptor library, 25 μL 2×KAPA mix, 2.5 μL I5 primer, 2.5 μL I7 primer, and 15 μL water were mixed in PCR tubes and tubes were incubated in a thermocycler. The program was set as 98°C 45 sec, (98°C 15 sec, 60°C 10 sec) ×14 cycles, 4°C hold. The library was purified using 1.4×DNA beads (Vazyme) and eluted in 25 μL water. The concentration of library was measured using qubit.

### CUT&Tag

CUT&Tag experiments were performed as previously described with modifications ^18^. Briefly, cells suspended in washing buffer were Concanavalin A coated magnetic beads (Polyscience, 86057) for 10 min at room temperature. Cells were then incubated with primary antibody (H3K4me3, Active motif #39160; H3K27me3, Cell Signaling Technology 9733 C36B11; H2AK119ub, Cell Signaling Technology 8240s; H3K9me3, active motif #39161) at a ratio of 1:100 at 4°C for 2 h on a thermomixer at 400 rpm. After the removal of primary antibody, secondary antibody (Invitrogen, #SA5-10228) were added at a ratio of 1:100, and incubation was performed for 1 h at 4°C. After washing two times, beads were incubated with protein A-Tn5 (Vazyme, S603) at 4°C for 1 h. Tagmentation was performed at 37°C for 30 min after washing. SDS was then added to stripe Tn5 and tagmented DNA was purified with 3 × VAHTS DNA Clean Beads (Vazyme, N411). Libraries were then performed by PCR with a library preparation kit (Vazyme, TD501) and sequencing was done using the Novaseq system (Illumina) according to the manufacturer’s protocol.

### sisHiC

sisHiC experiments were performed as previously described ^36^.

### RNA-seq

RNA-seq libraries were prepared using Smart-seq2 method as previously reported ^37^.

### ChIP-seq and CUT&Tag data analysis

The paired end ChIP-seq or CUT&Tag reads were mapped to mm9 using bowtie2 (version 2.2.5) with the parameters: -t -q -p 40 -N 1 -L 25 -X 2000 --no-mixed --no-discordant. The unmapped reads, not uniquely mapped, and duplicate reads were removed. Read coverages over the mm9 genome were calculated by bamCoverage from deepTools v3.3.1. The signal was then normalized by computing the number of reads per kilobase of bin per million reads sequenced (RPKM), followed by normalization using Z-score (whole genome 100bp bn with outliner regions excluded). The signal was then visualized using UCSC genome browser. To calculate the average ChIP-seq signal on gene desert or gene forest regions, these two regions were first defined as previously described ^36^. Each region was then split to 10 bins and ChIP-seq or CUT&Tag signal were calculated to these bins and average signal in each bin was then calculated. To calculate and plot the correlation between H3K9me3 and CBX7, the 100bp bin, Z-score normalized signals were then re-calculated to 2000 bp bin and the 2000 bp bin signal of CBX7 ChIP-seq and H3K9me3 CUT&Tag were merged to plot scatter plot and calculate pearson correlation.

### Hi-C data analysis

Hi-C data analysis was performed as previously described ^36^. Briefly, Hi-C libraries were aligned using HiCPro (version 2.10.0) with bowtie2 end-to-end algorithm and with the option ‘-very-sensitive’. The validpairs files were converted to hic files using HiCPro and then visualized by juicebox to generate Hi-C heatmaps. To calculate the number of compartments of each chromosome in each sample, the validpairs files were converted to mcool files using cooltools. Eigenvectors were calculated by cooltools (Open2C et al., 2022) and compartment values were extracted from the E1 column of the eigenvectors.

### RNA-seq analysis

Paired-end RNA-seq reads were aligned to mm9 using HISAT2 v2.2.1 and FPKM value of each gene was calculated with StringTie v2.1.2. HTSeq v0.6.0 was used to calculate the counts of each gene. Differentially expressed genes (DEGs) were calculated with DESeq2 v1.24.0 with the cutoff pvalue < 0.01 and foldchange > 2. Gene ontology analysis was performed with gseGO function in clusterProfiler package in R.

### Data availability

The raw sequencing data and processed data of this paper (ChIP-seq/RNA-seq/CUT&Tag/Hi-C) have been deposited in the Gene Expression Omnibus database under the accession number: GSE241262 https://www.ncbi.nlm.nih.gov/geo/query/acc.cgi?acc=GSE241262

## SUPPLEMENTAL INFORMATION

Supplemental Information includes 3 tables.

## Acknowledgments

This work was supported by grants from National Key R&D Program (2019YFA0508403 to P.L. & 2019YFA0508900 to W.X.) and Natural Science Foundation of China (32330024, 32150023, 32125010 to P.L.; 31988101, 31830047, 31725018 to W.X.). The Tsinghua-Peking Center for Life Sciences (P.L. and W.X.). W.X. is a recipient of an HHMI International Research Scholar award and New Cornerstone Investigator. We thank Jun Chen and Cuifang Liu from Guohong Li’s lab at the Institute of Biophysics, Chinese Academy of Sciences for sharing plasmids of Xenopus laevis histone; Professor Bing Liu at the Institute of Hematology, Fifth Medical Center of Chinese PLA General Hospital for core blood donor; We thank Dr. Bofeng Liu, Dr. Zhenhai Du, and Dr. Fengling Chen in the Xie lab for suggestions in Hi-C, ChIP-seq, and RNA-seq analysis; We thank Dr. Ge Zhan from Xiaohua Shen’s lab at Tsinghua university for sharing probes; We thank Tsinghua Nikon Imaging Center for technical assistance, and other members in the Li Lab for discussion.

## Author contributions

P.L. conceived the idea; W.S. and J.W. carried out the experiments and analyzed the data; W.L. carried out the ITC assay and analyzed the data; B.T. provided technical support for CD34+ cord blood isolation; H.Q. captured electron microscopy images of nucleosomal arrays; H.L. and J.W. provided important technical guidance; P.L. and W.X. obtained funding and supervised the project; All authors participated in manuscript writing.

## Declaration of interests

P.L. is the scientific co-founder of NuPhase Therapeutics, whose projects are not related with this work. Other authors declare that they have no other conflict of interest.

---

**Extended Data Fig. 1.**
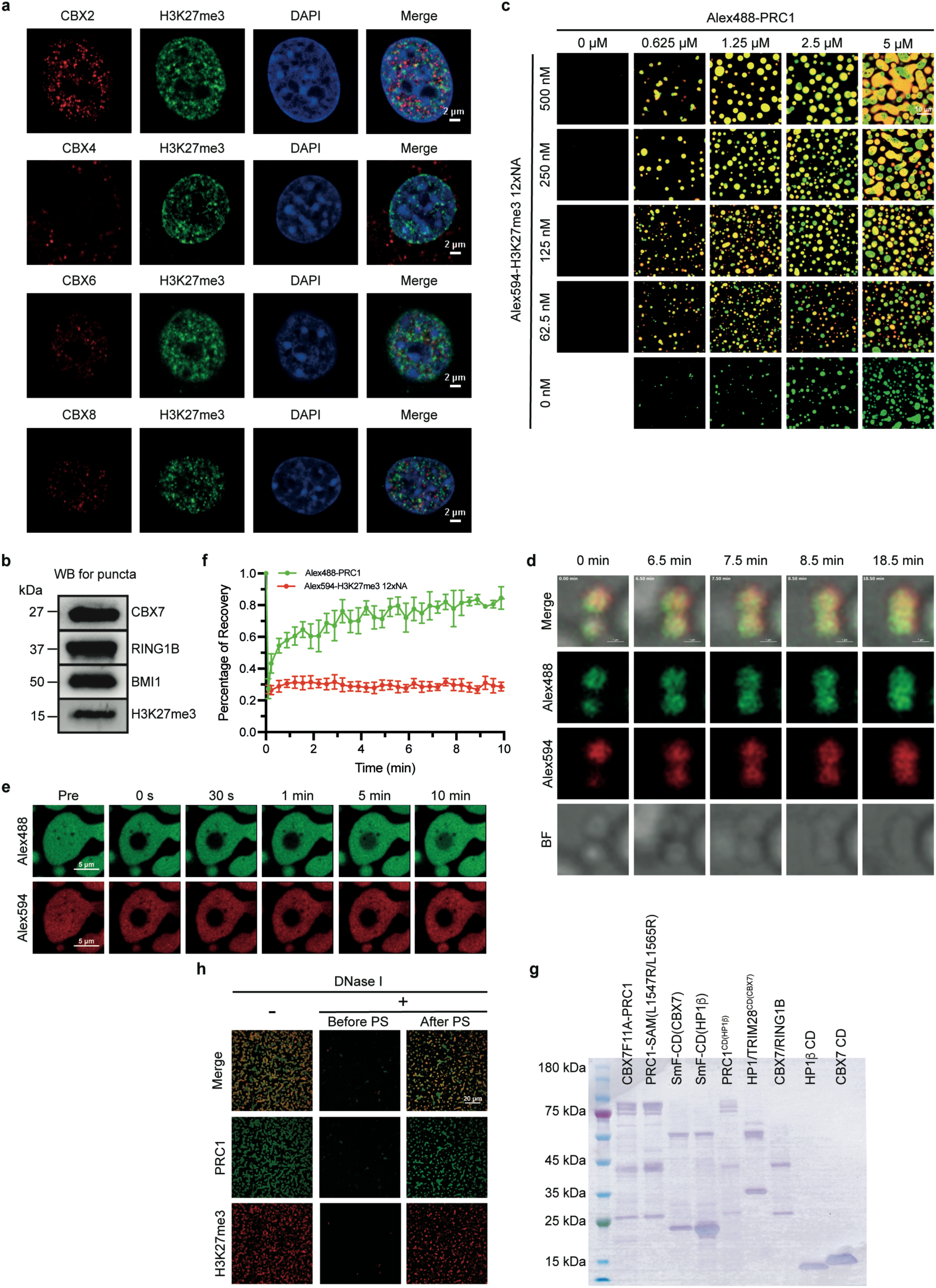
Multivalent H3K27me3-PRC1 interactions *in vivo* and *in vitro.* **a**, Immunofluorescence images of NIH-3T3 cells show that CBX2/4/6/8 (red) have less colocalization with H3K27me3 (green) than CBX7 in facultative heterochromatin. DNA was counterstained with DAPI (blue). **b**, Western blotting confirms that H3K27me3 is enriched in the puncta formed by PRC1 complex with H3K27me3-12xNA in Fig. 1l. **c,** Phase diagram of PRC1 complex with H3K27me3 12xNA. PRC1 complex was partially (5%) labeled with Alexa Fluor 488 (green), and histone 2B (H2B) was totally labeled with Alexa Fluor 594 (red). The images shown are for merged channels. Scale bar =10 μm. **d**, Droplets formed by LLPS of PRC1 (green) with H3K27me3 (red) fuse upon contact. Scale bar = 1 μm. **e**, Snapshots of a bleached droplet of PRC1 complex with H3K27me3 12xNA. PRC1 complex was partially (5%) labeled with Alexa Fluor 488 (green), and histone 2B (H2B) was totally labeled with Alexa Fluor 594 (red). Images show Alexa Fluor 488 and Alexa Fluor 594 signals. **f**, Quantification of average fluorescence intensity with time in the bleached region of droplets (n=3). All of the data are presented as mean ± SDs. **g**, SDS-PAGE analysis of the proteins in this paper. **h**, The droplets resulting from LLPS of H3K27me3 12xNA with PRC1 complex are resistant to DNase I. H3K27me3 12xNA was treated by buffer (left) or DNase I (center) before being mixed with PRC1 complex. H3K27me3 12xNA was mixed with PRC1 before DNase I treatment. PS, phase separation. PRC1 concentration, 5 μM; H3K27me3 12xNA concentration, 250 nM. Scale bar: 20 μm.

**Extended Data Fig. 2.**
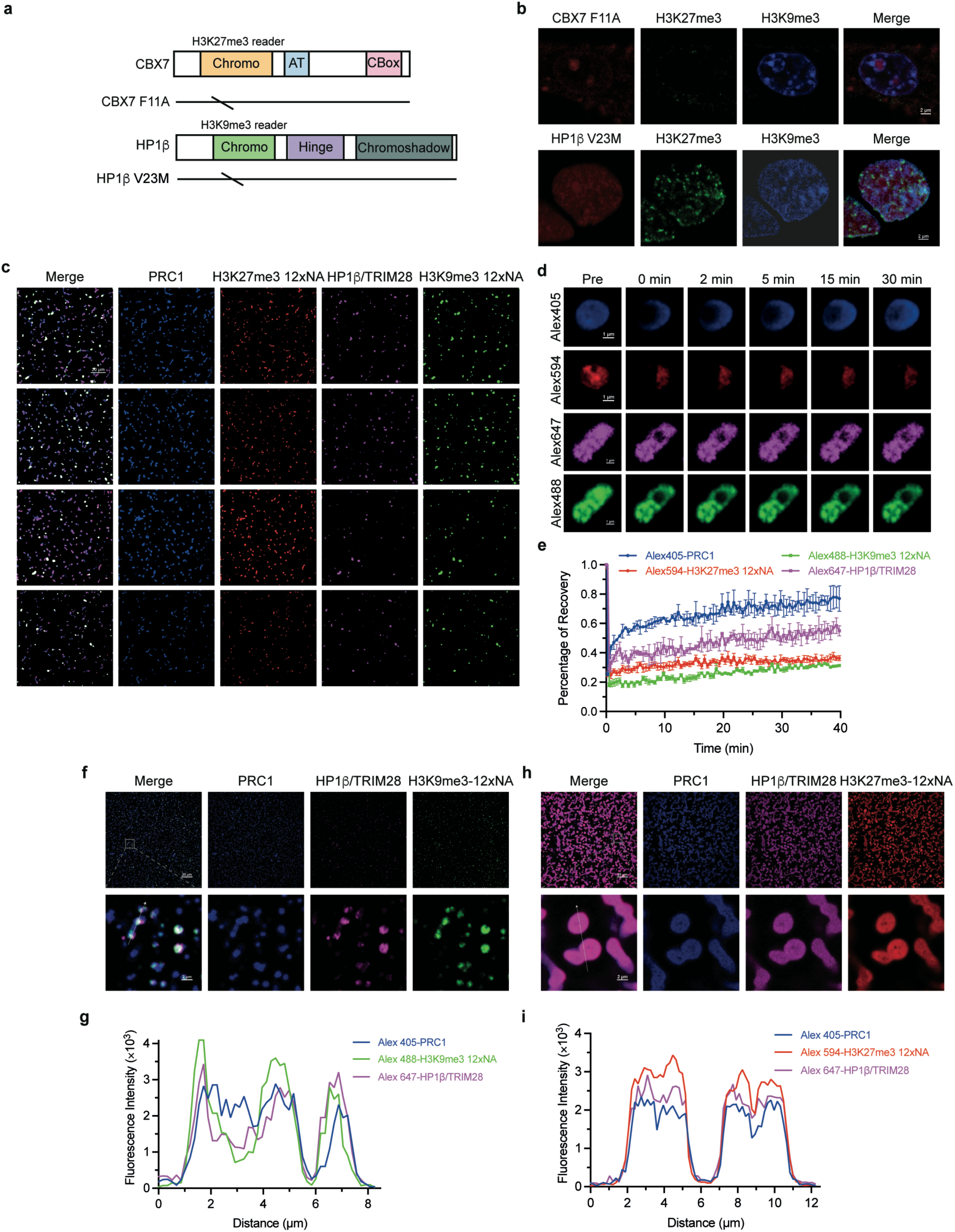
LLPS-mediated compartmentalization is driven by multivalent PTM-reader interactions. **a**, Domain structures and mutated forms of CBX7 and HP1β. **b**, The number of facultative heterochromatin puncta is reduced upon overexpression of mCherry-CBX7(F11A) in NIH-3T3 cells. The number of constitutive heterochromatin puncta is reduced upon overexpression of mCherry-HP1β(V23M) in NIH-3T3 cells. **c**, The images show that H3K27me3-PRC1 droplets are immiscible with H3K9me3-HP1β/TRIM28 droplets at four different concentrations *in vitro*. Scale bar = 20 μm. **d**, Snapshots of bleached droplets of PRC1 with H3K27me3 12xNA and HP1β/TRIM28 with H3K9me3 12xNA. **e**, Quantification of average fluorescence intensity with time in the bleached region of droplets (n=3). All of the data are presented as means ± SDs. **f**, Images of PRC1 mixed with H3K9me3-HP1β/TRIM28 droplets. The gray box indicates a zoomed-in area shown at the bottom. Scale bar = 2 μm. **g**, Line scans of the images of droplets at the position depicted by the white arrow in the merged image of the bottom row in (**f**). **h**, Images of HP1β/TRIM28 mixed with H3K27me3-PRC1 droplets. The gray box indicates a zoomed-in area shown at the bottom. Scale bar = 2 μm. **i**, Line scans of the images of droplets at the position depicted by the white arrow in the merged image of the bottom row in (**h**).

**Extended Data Fig. 3.**
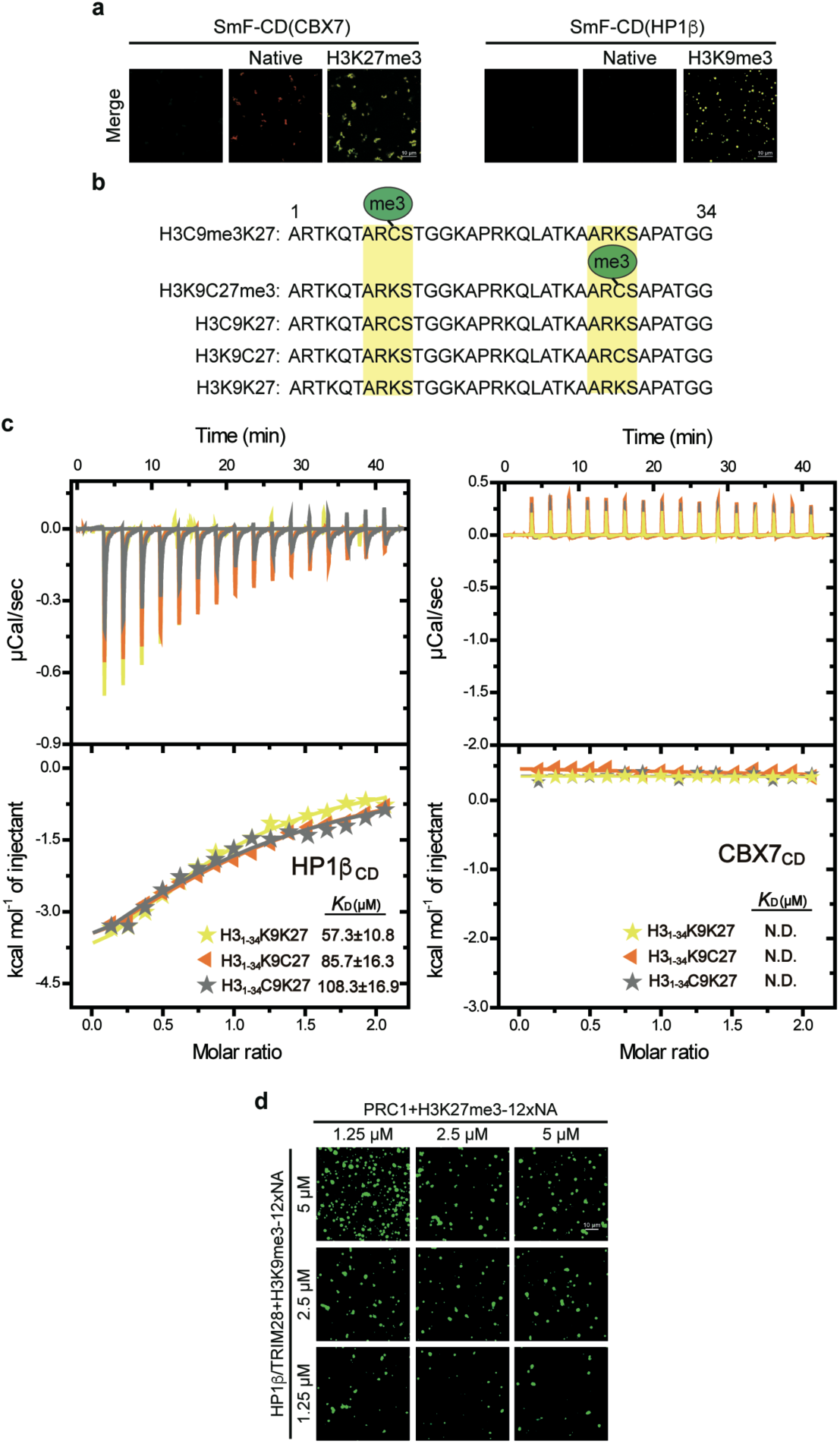
The mechanism underlying immiscibility. **a**, Left: Images of SmF-CD(CBX7) alone, mixed with unmodified 12xNA, or H3K27me3 12xNA. SmF-CD(CBX7) was partially (5%) labeled with Alexa Fluor 488 (green), H2B was totally labeled with Alexa Fluor 594 (red) in unmodified and H3K27me3 12xNA. SmF-CD(CBX7), 20 μM; 12xNA, 2 μM. Scale bar = 10 μm. Right: Images of SmF-CD(HP1β) alone, mixed with unmodified 12xNA, or H3K9me3 12xNA. SmF-CD(HP1β) was partially (5%) labeled with Alexa Fluor 594 (red), H2B was totally labeled with Alexa Fluor 488 (green) in unmodified and H3K9me3 12xNA. SmF-CD(HP1β), 5 μM; 12xNA, 500 nM. The images shown are for merged channels. Scale bar=10 μm. **b**, Amino acid sequences of modified and unmodified histone 3 peptides in ITC assays. **c**, ITC curves of H3C9K27, H3K9C27, and H3K9K27 peptides titrated into HP1β and CBX7 CD. N.D., not detected. **d**, In vitro images of H3K27me3-PRC1 mixed with H3K9me3-HP1β/TRIM28 droplets. The green signal shows the occupation of H3K9me3. PRC1 concentrations are labeled, and H3K27me3 12xNA concentrations are one-tenth of PRC1. HP1β/TRIM28 concentrations are labeled, and H3K9me3 12xNA concentrations are one-fifth of HP1β/TRIM28. Scale bar = 10 μm.

**Extended Data Fig. 4.**
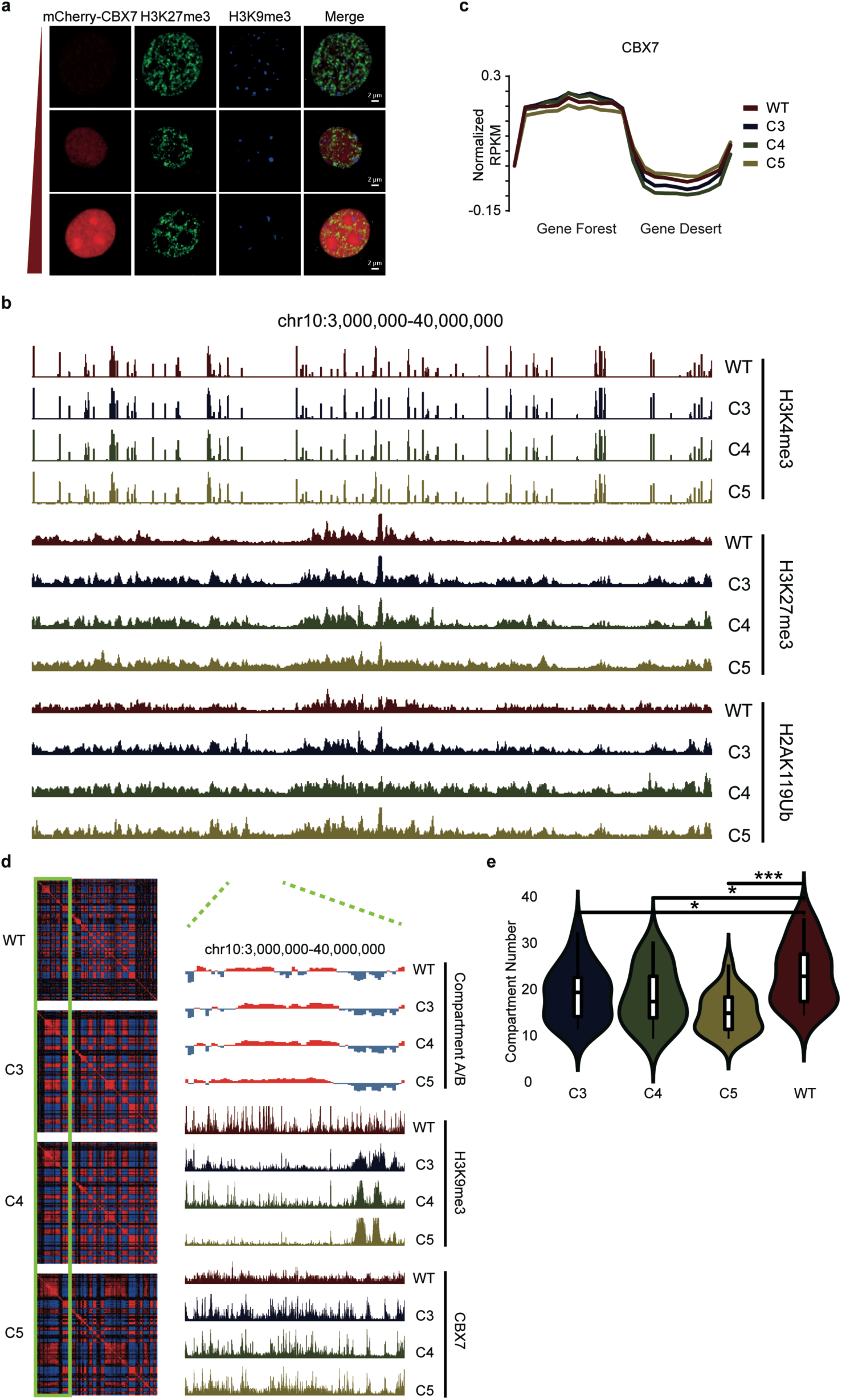
PRC1 over-expression disrupts the chromatin higher order structure. **a**, Confocal fluorescence images of H3K27me3 (green) and H3K9me3 (blue) in NIH-3T3 cell lines with the stable expression of mCherry-CBX7, cells with CBX7 expression from low (top row) to high (bottom row). Scale bar = 2 μm. **b**, UCSC browser view showing H3K4me3, H3K27me3, and H2AK119ub were not affected in mCherry-CBX7 clones. **c**, Average CBX7 enrichment (Z-score normalized RPKM) at gene desert region and gene forest region in wild-type cells and mCherry-CBX7 clones. **d**, Heatmaps of correlation matrix (left) showing the global change of genome contact and UCSC browser views in mCherry-CBX7 clones. Two dashed green lines showing the change of compartment A/B and deposition of H3K9me3 in the clones are correlated with CBX7 binding in the region in the green box. **e**, Violin plot and box plot showing the distribution of the number of compartment B in each chromosome in mCherry-CBX7 clones or WT cells. Representative *p* values are for student *t*-test between WT cells and each CBX7 OE clone (*, *p*<0.1; ***, *p*<0.001).

**Extended Data Fig. 5.**
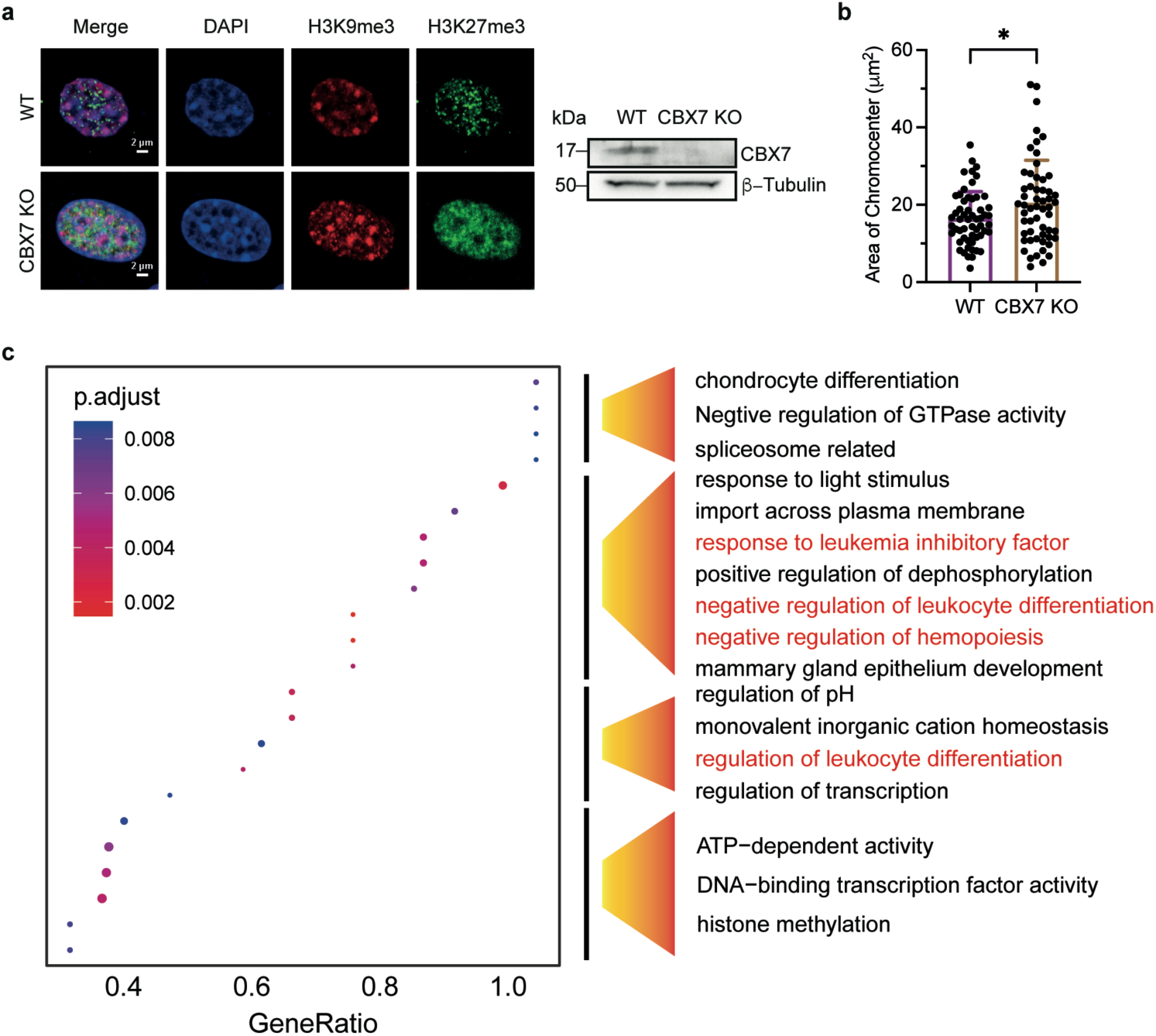
Overexpression of CBX7 causes alteration of epigenetic patterns and is related to the emergence of AML. **a**, Left: Immunofluorescence images of wild-type and CBX7 knock-out NIH-3T3 cells. DNA was counterstained with DAPI (blue). Right: Western blot of CBX7 in wild-type and CBX7 knock-out cells. **b**, Area of chromocenters in wild-type 3T3 cells compared to CBX7 knock-out cells. *p* values were calculated using a two-sided unpaired student *t*-test (****, *p*<0.0001). All of the data are presented as means+SDs. **c**, GO analysis of the top 250 genes that loss of H3K9me3 mark in the mCherry-CBX7 clones, *p* value < 0.01, AML-related terms are labeled red.

**Extended Data Fig. 6.**
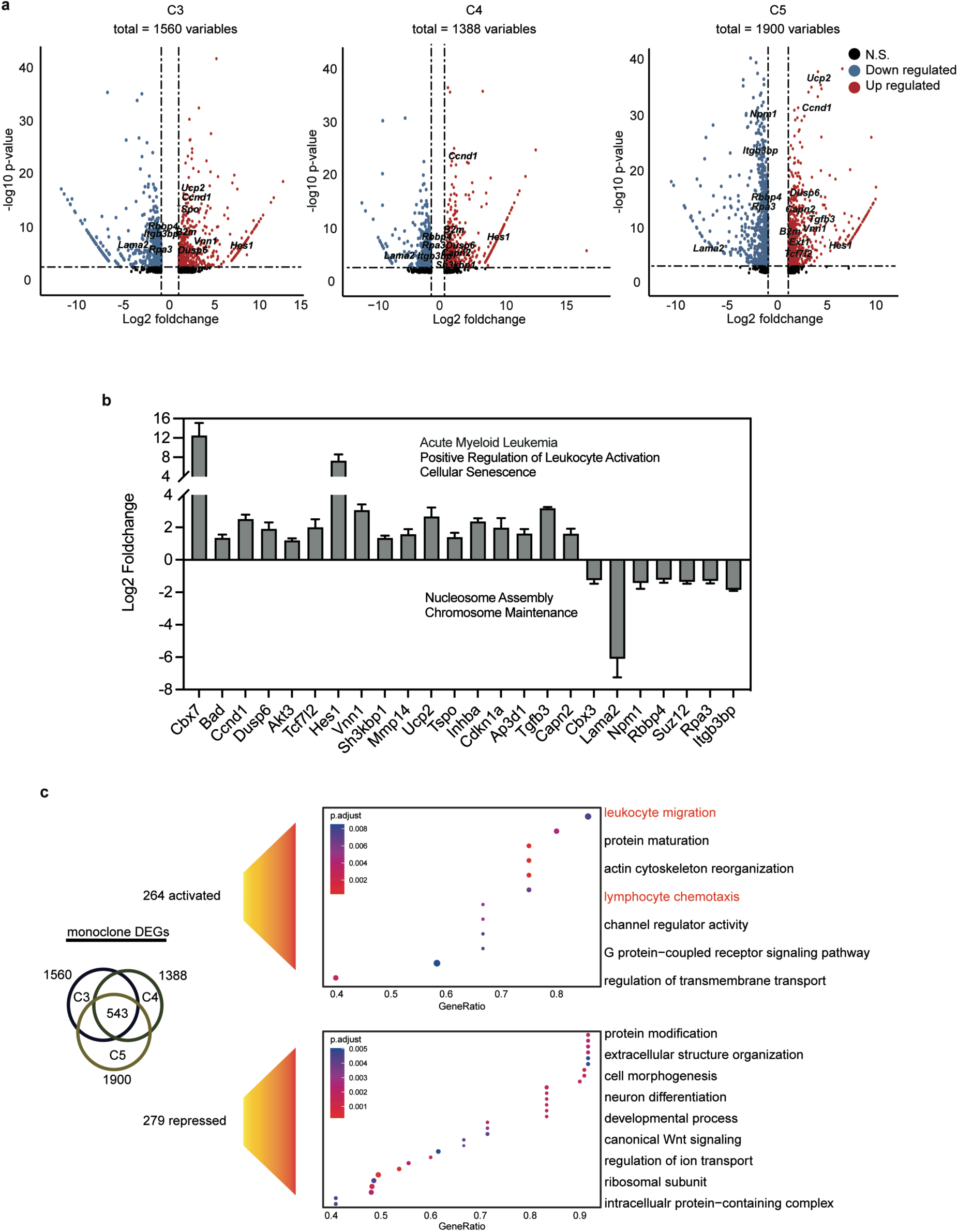
Overexpression of CBX7 causes the emergence of AML. **a**, Volcano plot showing the up-regulated genes (red) and down-regulated genes (blue) in mCherry-CBX7 clone 3, clone 4, and clone 5 with cut-off of log2 foldchange > 1 and *p* value < 0.01. **b**, Bar plot showing the log2 foldchange differentially expressed genes related to AML and chromatin organization. **c**, Vein plot showing the number of differentially expressed genes in mCherry-CBX7 clones and GO analysis of 543 overlapped genes among the clones, *p* value < 0.01, AML-related terms are labeled in red.

**Extended Data Fig. 7.**
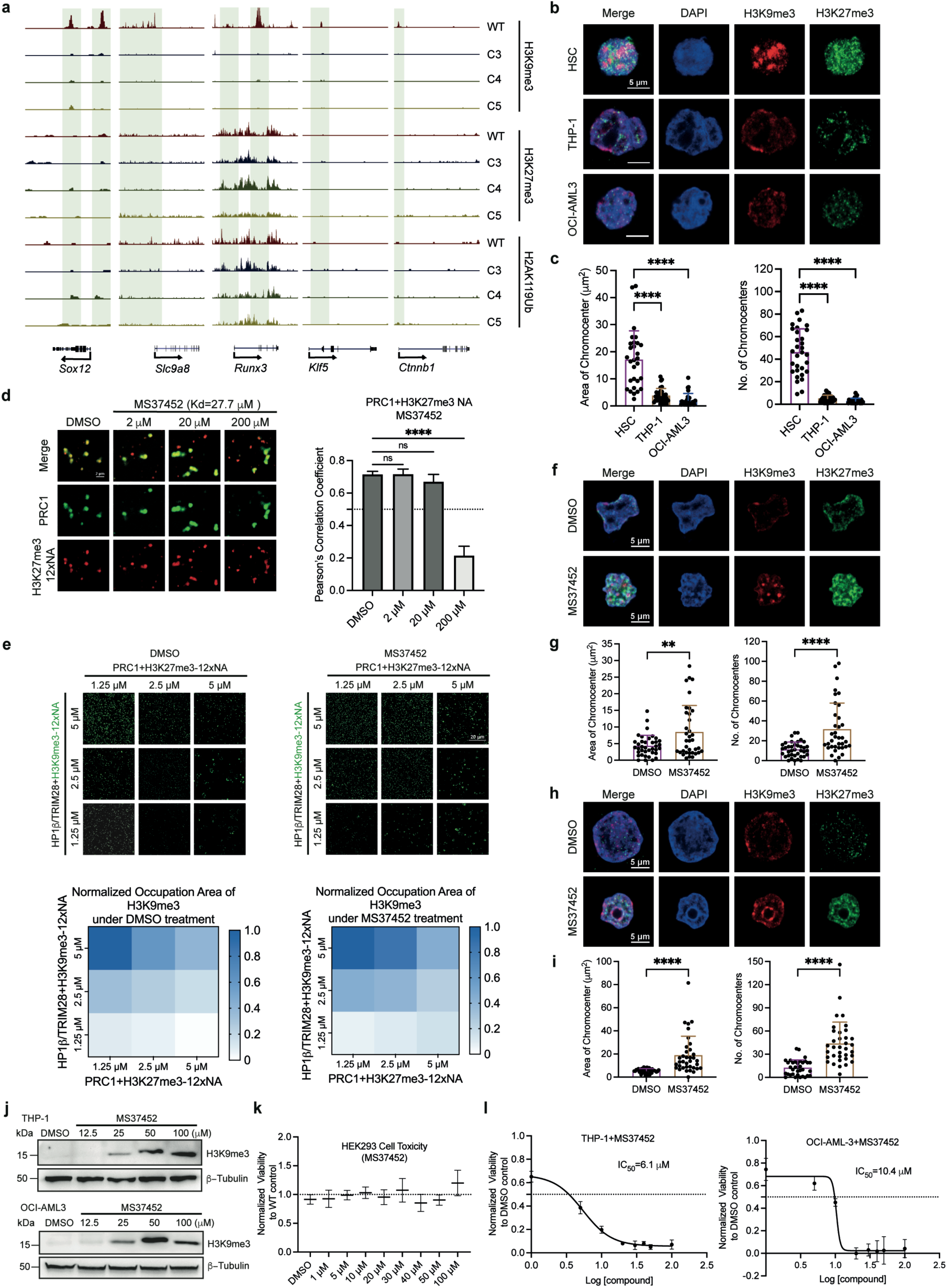
Inhibition of CBX7-PRC1 phase separation results in the reorganization of H3K9me3. **a**, UCSC browser view showing the loss of H3K9me3 at the promoters of AML-related genes in mCherry-CBX7 clones. No dramatic changes in H3K27me3 and H2AK119Ub at the promoters. **b**, Immunofluorescence images of H3K9me3 (red) and H3K27me3 (green) in CD34^+^ HSCs and two AML cell lines, THP-1 and OCI-AML3. Nuclei were stained by DAPI (blue). **c**, Area and number of chromocenters in HSC, THP-1, and OCI-AML3 cells. Representative *p* values are for student *t*-test (****, *p*<0.0001). **d**, Phase-separated fluorescence images of PRC1 (green) with reconstituted H3K27me3 12xNA (red) under the conditions of DMSO and MS37452. Scale bar = 2 μm. MS37452 blocks the interaction between CBX7-PRC1 and H3K27me3 12xNA, as determined by the Pearson’s correlation coefficients of green and red signals. Data are presented as means+SDs and *p* values (calculated by student *t*-test) are for comparison to the DMSO control (*, *p*<0.0332; **, *p*<0.0021; ****, *p*<0.0001; ns, not significant). **e**, H3K9me3 compartment can be partially reconstructed after MS37452 (100 μM) treatment *in vitro*. A heat map showing the normalized area occupied by H3K9me3 signals in the H3K9me3-HP1β/TRIM28 and H3K27me3-PRC1 mixing assays. Scale bar = 20 μm. **f**, Immunofluorescence images of H3K9me3 (red) and H3K27me3 (green) in THP-1 cells after 6 days of treatment with DMSO or 100 μM MS37452. Nuclei were stained by DAPI (blue). Scale bar = 5 μm. **g**, Area and number of chromocenters in THP-1 cells incubated MS37452 or DMSO. **h**, Immunofluorescence images of H3K9me3 (red) and H3K27me3 (green) in OCI-AML3 cells after 6 days of treatment with DMSO or 100 μM MS37452. Nuclei were stained by DAPI (blue). Scale bar = 5 μm. **i**, Area and number of chromocenters in OCI-AML3 cells incubated MS37452 or DMSO. In (**g**) and (**i**), data are presented as means+SDs and *p* values are calculated by student *t*-test (**, *p*<0.0021; ****, *p*<0.0001). **j**, Western blot analysis of H3K9me3 in THP1 and OCI-AML3 cells under MS37452 treatment. **k**, MS37452 displays no toxicity up to 100 μM in HEK293 cells. Data are presented as means ± SDs. **l**, MS37452 inhibits the proliferation of THP-1 and OCI-AML3 cells in a dose-dependent manner. Cells were incubated with inhibitor for 6 days (n = 5, two biological replicates with five technical replicates). Data are presented as means ± SDs.

## Declare of interests

P.L. is the scientific co-founder of NuPhase Therapeutics, whose projects are not related with this work. Other authors declare that they have no conflict of interest.

## References

1 Wang, L. et al. Histone Modifications Regulate Chromatin Compartmentalization by Contributing to a Phase Separation Mechanism. Mol Cell 76, 646–659 e646, doi:10.1016/j.molcel.2019.08.019 (2019).

2 Fischle, W. et al. Molecular basis for the discrimination of repressive methyl-lysine marks in histone H3 by Polycomb and HP1 chromodomains. Genes Dev 17, 1870–1881, doi:10.1101/gad.1110503 (2003).

3 Gibson, B. A. et al. Organization of Chromatin by Intrinsic and Regulated Phase Separation. Cell 179, 470–484 e421, doi:10.1016/j.cell.2019.08.037 (2019).

4 Ruthenburg, A. J., Li, H., Patel, D. J. & Allis, C. D. Multivalent engagement of chromatin modifications by linked binding modules. Nat Rev Mol Cell Biol 8, 983–994, doi:10.1038/nrm2298 (2007).

5 Jenuwein, T. & Allis, C. D. Translating the histone code. Science 293, 1074–1080, doi:10.1126/science.1063127 (2001).

6 Schuettengruber, B., Bourbon, H. M., Di Croce, L. & Cavalli, G. Genome Regulation by Polycomb and Trithorax: 70 Years and Counting. Cell 171, 34–57, doi:10.1016/j.cell.2017.08.002 (2017).

7 Blackledge, N. P., Rose, N. R. & Klose, R. J. Targeting Polycomb systems to regulate gene expression: modifications to a complex story. Nat Rev Mol Cell Biol 16, 643–649, doi:10.1038/nrm4067 (2015).

8 Seif, E. et al. Phase separation by the polyhomeotic sterile alpha motif compartmentalizes Polycomb Group proteins and enhances their activity. Nat Commun 11, 5609, doi:10.1038/s41467-020-19435-z (2020).

9 Simon, M. D. et al. The site-specific installation of methyl-lysine analogs into recombinant histones. Cell 128, 1003–1012, doi:10.1016/j.cell.2006.12.041 (2007).

10 Bernstein, E. et al. Mouse polycomb proteins bind differentially to methylated histone H3 and RNA and are enriched in facultative heterochromatin. Mol Cell Biol 26, 2560–2569, doi:10.1128/MCB.26.7.2560-2569.2006 (2006).

11 Robinson, A. K. et al. The growth-suppressive function of the polycomb group protein polyhomeotic is mediated by polymerization of its sterile alpha motif (SAM) domain. J Biol Chem 287, 8702–8713, doi:10.1074/jbc.M111.336115 (2012).

12 Kim, C. A., Gingery, M., Pilpa, R. M. & Bowie, J. U. The SAM domain of polyhomeotic forms a helical polymer. Nat Struct Biol 9, 453–457, doi:10.1038/nsb802 (2002).

13 Nichols, M. H. & Corces, V. G. Principles of 3D compartmentalization of the human genome. Cell Rep 35, 109330, doi:10.1016/j.celrep.2021.109330 (2021).

14 Collins, B. M. et al. Homomeric ring assemblies of eukaryotic Sm proteins have affinity for both RNA and DNA. Crystal structure of an oligomeric complex of yeast SmF. J Biol Chem 278, 17291–17298, doi:10.1074/jbc.M211826200 (2003).

15 Zhou, M. et al. Phase-separated condensate-aided enrichment of biomolecular interactions for high-throughput drug screening in test tubes. J Biol Chem 295, 11420–11434, doi:10.1074/jbc.RA120.012981 (2020).

16 Rowley, M. J. & Corces, V. G. Organizational principles of 3D genome architecture. Nat Rev Genet 19, 789–800, doi:10.1038/s41576-018-0060-8 (2018).

17 Lieberman-Aiden, E. et al. Comprehensive mapping of long-range interactions reveals folding principles of the human genome. Science 326, 289–293, doi:10.1126/science.1181369 (2009).

18 Kaya-Okur, H. S. et al. CUT&Tag for efficient epigenomic profiling of small samples and single cells. Nat Commun 10, 1930, doi:10.1038/s41467-019-09982-5 (2019).

19 Klauke, K. et al. Polycomb Cbx family members mediate the balance between haematopoietic stem cell self-renewal and differentiation. Nat Cell Biol 15, 353–362, doi:10.1038/ncb2701 (2013).

20 Jung, J. et al. CBX7 Induces Self-Renewal of Human Normal and Malignant Hematopoietic Stem and Progenitor Cells by Canonical and Non-canonical Interactions. Cell Rep 26, 1906–1918 e1908, doi:10.1016/j.celrep.2019.01.050 (2019).

21 Verhaak, R. G. et al. Prediction of molecular subtypes in acute myeloid leukemia based on gene expression profiling. Haematologica 94, 131–134, doi:10.3324/haematol.13299 (2009).

22 Gang, D. et al. Aging-related genes related to the prognosis and the immune microenvironment of acute myeloid leukemia. Clin Transl Oncol, doi:10.1007/s12094-023-03168-8 (2023).

23 Wang, J., Wu, C. & Zhou, W. CircPLXNB2 regulates acute myeloid leukemia progression through miR-654-3p/CCND1 axis. Hematology 28, 2220522, doi:10.1080/16078454.2023.2220522 (2023).

24 Wang, H. et al. Disruption of dNTP homeostasis by ribonucleotide reductase hyperactivation overcomes AML differentiation blockade. Blood 139, 3752–3770, doi:10.1182/blood.2021015108 (2022).

25 Kato, T. et al. Hes1 suppresses acute myeloid leukemia development through FLT3 repression. Leukemia 29, 576–585, doi:10.1038/leu.2014.281 (2015).

26 Brown, J., Greaves, M. F. & Molgaard, H. V. The gene encoding the stem cell antigen, CD34, is conserved in mouse and expressed in haemopoietic progenitor cell lines, brain, and embryonic fibroblasts. Int Immunol 3, 175–184, doi:10.1093/intimm/3.2.175 (1991).

27 Ren, C. et al. Small-molecule modulators of methyl-lysine binding for the CBX7 chromodomain. Chem Biol 22, 161–168, doi:10.1016/j.chembiol.2014.11.021 (2015).

28 Moller, M. et al. Destabilization of chromosome structure by histone H3 lysine 27 methylation. PLoS Genet 15, e1008093, doi:10.1371/journal.pgen.1008093 (2019).

29 Peters, A. H. et al. Partitioning and plasticity of repressive histone methylation states in mammalian chromatin. Mol Cell 12, 1577–1589, doi:10.1016/s1097-2765(03)00477-5 (2003).

30 Cooper, S. et al. Targeting polycomb to pericentric heterochromatin in embryonic stem cells reveals a role for H2AK119u1 in PRC2 recruitment. Cell Rep 7, 1456–1470, doi:10.1016/j.celrep.2014.04.012 (2014).

31 Gil, J., Bernard, D., Martinez, D. & Beach, D. Polycomb CBX7 has a unifying role in cellular lifespan. Nat Cell Biol 6, 67–72, doi:10.1038/ncb1077 (2004).

32 Stevens, T. J. et al. 3D structures of individual mammalian genomes studied by single-cell Hi-C. Nature 544, 59–64, doi:10.1038/nature21429 (2017).

33 Luger, K., Rechsteiner, T. J. & Richmond, T. J. Expression and purification of recombinant histones and nucleosome reconstitution. Methods Mol Biol 119, 1–16, doi:10.1385/1-59259-681-9:1 (1999).

34 Lowary, P. T. & Widom, J. New DNA sequence rules for high affinity binding to histone octamer and sequence-directed nucleosome positioning. J Mol Biol 276, 19–42, doi:10.1006/jmbi.1997.1494 (1998).

35 Zhang, Y. et al. Transcriptionally active HERV-H retrotransposons demarcate topologically associating domains in human pluripotent stem cells. Nat Genet 51, 1380–1388, doi:10.1038/s41588-019-0479-7 (2019).

36 Du, Z. et al. Allelic reprogramming of 3D chromatin architecture during early mammalian development. Nature 547, 232–235, doi:10.1038/nature23263 (2017).

37 Picelli, S. et al. Full-length RNA-seq from single cells using Smart-seq2. Nat Protoc 9, 171–181, doi:10.1038/nprot.2014.006 (2014).

38 Gray, F. et al. BMI1 regulates PRC1 architecture and activity through homo- and hetero-oligomerization. Nat Commun 7, 13343, doi:10.1038/ncomms13343 (2016).

