## Supplementary material for "Histone marks enable formation of immiscible phase-separated chromatin compartments": Table S1-S3: Supplementary Information.docx

Table S1. ITC raw data.

|  | **Peptide** | **ΔH（cal/mol）** | **ΔS（cal/mol/deg）** | **K_a_（M^-1^）** | **N** | **Chi^2/DoF** |
| --- | --- | --- | --- | --- | --- | --- |
|  | H3_1-34_C9me3 | -8523±70.29 | -2.47 | 5.09E5±4.30E4 | 1.11±0.00652 | 1.632E4 |
|  | H3_1-34_C27me3 | -6076±513.0 | -1.30 | 1.48E4±1.94E3 | 0.960±0.0518 | 2889 |
| HP1_CD_ | H3_1-34_K9K27 | -5982±695.4 | -0.578 | 1.81E4±3.41E3 | 0.871±0.0675 | 7722 |
|  | H3_1-34_K9C27 | -6051±736.1 | -1.61 | 1.21E4±2.30E3 | 1.10±0.0794 | 4366 |
|  | H3_1-34_C9K27 | -7090±482.9 | -5.58 | 9.47E3±1.48E3 | 1.00±0 | 1.679E4 |
|  | H3_1-34_C9me3 | N.D. | | | | |
|  | H3_1-34_C27me3 | N.D. | | | | |
| CBX7_CD_ | H3_1-34_K9K27 |  |  | N.D. |  | |
|  | H3_1-34_K9C27 | -925.3±43.27 | 19.2 | 7.34E4±1.82E4 | 0.852±0.0292 | 1553 |
|  | H3_1-34_C9K27 | -603.8±22.22 | 20.7 | 9.09E4±2.24E4 | 1.08±0.0290 | 773.3 |

Table S2. Source of antibodies.

| **Antibodies** | **Source** | **ID** |
| --- | --- | --- |
| Histone H3K4me3 antibody (pAb) | Active Motif | Cat# 39159; RRID: AB_2615077 |
| Rabbit Anti-Histone H3, trimethyl(Lys9) | Abcam | Cat# ab8898; RRID: AB_306848 |
| Histone H3K9me3 antibody (pAb) | Active Motif | Cat# 39161 |
| Tri-Methyl-Histone H3 (Lys27), Rabbit mAb | Cell signaling technology | Cat# 9733T |
| Goat anti-Rabbit IgG Fc Secondary Antibody | Invitrogen | #SA5-10228 |
| Anti-Histone H3 (tri methyl K27) | Abcam | Cat# ab6002 |
| Ubiquityl-Histone H2A (Lys119) Rabbit mAb | Cell signaling technology | Cat# 8240S |
| HP1α antibody | Abcam | Cat# ab77256; RRID: AB_1523784 |
| HP1β antibody | Abcam | Cat# ab10811; RRID: AB_297490 |
| HP1γ antibody | Abcam | Cat# ab10480; RRID: AB_297219 |
| Anti-CBX2 Mouse Monoclonal Antibody | HUABIO | Cat# EM1706-28 |
| Anti-Cbx4 antibody | Abcam | Cat# ab139815 |
| Anti-Cbx6 antibody | Abcam | Cat# ab229256 |
| Anti-Cbx7 antibody | Abcam | Cat# ab21873 |
| CBX8 Polyclonal Antibody | Invitrogen | Cat# PA5-103359 |
| Anti-RING2 / RING1B / RNF2 Antibody | Abcam | Cat# ab101273 |
| Anti-BMI1 Antibody | Abcam | Cat# ab85688 |
| Anti-PHC1 Antibody | Abcam | Cat# ab175424 |
| Lamin B1 Rabbit Polyclonal Antibody | Proteintech | Cat# 12987-1-AP |
| β-Tubulin Mouse mAb | ABclonal | Cat# AC021 |
| Donkey anti-Mouse IgG (H+L) Highly Cross-Adsorbed Secondary Antibody, Alexa Fluor™ 488 | Abcam | Cat# ab150105 |
| TRITC-Conjugated goat anti-Rat IgG antibody | ZSGB | Cat# ZF-0318 |
| Donkey anti-Rabbit IgG (H+L) Highly Cross-Adsorbed Secondary Antibody, Alexa Fluor™ 546 | Thermofisher Scientific | A10040 |
| Donkey anti-Mouse IgG (H+L) Highly Cross-Adsorbed Secondary Antibody, Alexa Fluor™ 546 | Thermofisher Scientific | A10036 |
| Donkey Anti-Rabbit lgG H&L(Alexa Fluor 647) antibody | Abcam | Cat# Ab150063; RRID: AB_2687541 |

Table S3. Software and algorithms.

| Excel | This paper; | [https://products.office.com/ en-us/excel](https://products.office.com/%20en-us/excel) |
| --- | --- | --- |
| ImageJ | Schneider et al.^1^ | <https://imagej.nih.gov/> |
| NIS-Elements AR Analysis | This paper; | <http://www.nis-elements.cz/> |
| GraphPad Prism | This paper; | [https://www.graphpad.com/ scientific-software/prism/](https://www.graphpad.com/%20scientific-software/prism/) |
| Bowtie2 | Langmead and Salzberg^2^ | <http://bowtie-bio.sourceforge.net/bowtie2/index.shtml> |
| DeepTools | Ramirez et al. ^3^ | <https://github.com/deeptools/deepTools> |
| HiC-Pro | Servant et al. ^4^ | <https://github.com/nservant/HiC-Pro> |
| Juicebox | Durand et al. ^5^ | <https://github.com/aidenlab/Juicebox> |
| Cooltools | Open2C et al., 2022 | <https://github.com/open2c/cooltools> |
| HISAT2 | Kim et al. ^6^ | <http://daehwankimlab.github.io/hisat2/> |
| StringTie | Pertea et al. ^7^ | <https://github.com/gpertea/stringtie> |
| HTSeq | Anders et al. ^8^ | <https://github.com/htseq> |
| DESeq2 | Love et al. ^9^ | <https://github.com/mikelove/DESeq2> |

1	Schneider, C. A., Rasband, W. S. & Eliceiri, K. W. NIH Image to ImageJ: 25 years of image analysis. *Nat Methods* **9**, 671-675, doi:10.1038/nmeth.2089 (2012).

2	Langmead, B. & Salzberg, S. L. Fast gapped-read alignment with Bowtie 2. *Nat Methods* **9**, 357-359, doi:10.1038/nmeth.1923 (2012).

3	Ramirez, F. *et al.* deepTools2: a next generation web server for deep-sequencing data analysis. *Nucleic Acids Res* **44**, W160-165, doi:10.1093/nar/gkw257 (2016).

4	Servant, N. *et al.* HiC-Pro: an optimized and flexible pipeline for Hi-C data processing. *Genome Biol* **16**, 259, doi:10.1186/s13059-015-0831-x (2015).

5	Durand, N. C. *et al.* Juicebox Provides a Visualization System for Hi-C Contact Maps with Unlimited Zoom. *Cell Syst* **3**, 99-101, doi:10.1016/j.cels.2015.07.012 (2016).

6	Kim, D., Paggi, J. M., Park, C., Bennett, C. & Salzberg, S. L. Graph-based genome alignment and genotyping with HISAT2 and HISAT-genotype. *Nat Biotechnol* **37**, 907-915, doi:10.1038/s41587-019-0201-4 (2019).

7	Pertea, M. *et al.* StringTie enables improved reconstruction of a transcriptome from RNA-seq reads. *Nat Biotechnol* **33**, 290-295, doi:10.1038/nbt.3122 (2015).

8	Anders, S., Pyl, P. T. & Huber, W. HTSeq--a Python framework to work with high-throughput sequencing data. *Bioinformatics* **31**, 166-169, doi:10.1093/bioinformatics/btu638 (2015).

9	Love, M. I., Huber, W. & Anders, S. Moderated estimation of fold change and dispersion for RNA-seq data with DESeq2. *Genome Biol* **15**, 550, doi:10.1186/s13059-014-0550-8 (2014).
